# Neural and Behavioural Correlates of Variance of Sensory Evidence

**DOI:** 10.1101/2025.07.17.665271

**Authors:** Amir.M Mousavi-Harris, Jamal Esmaily, Germain Lefebvre, Joaquin Navajas, Bahador Bahrami

## Abstract

Neurobiology of perceptual decisions has largely focused on the neural correlates of the mean strength of sensory evidence. Much less is known about the neural coding of sensory variability. Here, we analyzed the EEG signals obtained from participants who judged the mean orientation of a sequence of gratings with varying variance but constant mean to identify the neural signatures of sensory variance and their relation to individual differences in choice confidence. The neural responses in the stimulus-entrained (4 Hz) and alpha (9–11 Hz) bands tracked variability independently of mean. The frontal and centro-parietal regions demonstrated a quadratic relationship (i.e., strongest responses to intermediate levels of uncertainty) to the standard deviation of the sequence. The occipital response coded the visual stimulus variability linearly. These neural markers of variability were correlated with inter-individual differences in computational components of metacognition. Centro-parietal activity was most predictive of metacognitive sensitivity, aligning with its known role in evidence accumulation. These findings advance our understanding of how the brain dynamically encodes uncertainty and help better characterise the electrophysiological basis of individual differences in metacognitive evaluation.

## Introduction

A large number of previous studies have examined the neuronal representation of sensory evidence in both animal and human brains, particularly through the lens of evidence accumulation and sequential sampling theories. Across animal studies, a fundamental finding is that the rate of evidence accumulation is proportional to the first moment of the evidence distribution, such as mean motion coherence, contrast, or stimulus amplitude. In non-human primates, single-unit recordings in the middle temporal (MT) area have shown that there are neurons in this area that encode motion strength, with firing rates that scale with the stimulus mean coherence, linking neural variability directly to behavioral fluctuations. Similarly, recordings in the lateral intraparietal (LIP) area have demonstrated that neural firing rates increase progressively at a rate proportion to mean motion coherence during decision-making tasks, with this rate of increase predicting both the speed and accuracy of choices, consistent with drift-diffusion models. These and similar findings in rodents support the tenet of sequential sampling models: decisions emerge as the integration of momentary evidence over time, with the accumulation rate tightly coupled to the average of the sensory input.

In humans, a similar relationship between evidence accumulation and the first moment of the evidence distribution has been confirmed using EEG and fMRI, with EEG studies providing the most direct neural signatures of sequential evidence integration. The work of Kelly and O’Connell (2013) (Kelly & O’Connell, 2013) has been particularly influential in establishing centroparietal positivity (CPP) as a robust index of evidence accumulation, demonstrating that its build-up rate is proportional to the mean coherence, contrast, or amplitude of sensory stimuli. Their studies show that stronger sensory evidence leads to steeper CPP slopes, reflecting faster accumulation and resulting in shorter reaction times, consistent with the predictions of drift-diffusion models. fMRI studies have similarly identified activity in the intraparietal sulcus (IPS) and dorsolateral prefrontal cortex (dlPFC) that scales with the strength of accumulated evidence. Furthermore, MEG studies have revealed that while accumulation is primarily driven by the mean strength of sensory input, it is also modulated by top-down influences such as prior expectations and task difficulty. Together, these findings provide strong support for the idea that human perceptual decision-making too operates under a sequential sampling framework, with the rate of evidence accumulation systematically reflecting the average of the sensory input.

In contrast to the extensive literature on the mean of sensory distributions, fewer studies have examined how higher-order statistical moments of sensory evidence, such as variance or standard deviation, are represented in the brain. Bang and Fleming (2018), Zylberberg et al. (2016), and Do et al. (2022) (Bang & Fleming, 2018; Do et al., 2022; Zylberberg et al., 2016) directly investigated neural responses to stimulus variance independent of the mean. Bang and

Fleming (2018) (Bang & Fleming, 2018) used human fMRI to examine how variance in sensory evidence affects the neural substrates of decision confidence, showing that higher variance led to greater uncertainty representations in the prefrontal cortex and parietal regions, independent of the mean strength of the sensory signals. Zylberberg et al. (2016) (Zylberberg et al., 2016), using monkey electrophysiology, found that neurons in the lateral intraparietal (LIP) cortex and the middle temporal (MT) area encoded stimulus variance, with higher variance reducing the rate of evidence accumulation and increasing decision variability. Do et al. (2022) (Do et al., 2022) employed human EEG to investigate sequential perceptual averaging, showing that higher stimulus variance affected the neural representation of accumulated sensory signals, particularly in the frontocentral cortex, reducing the precision of the mean representation over time.

While these studies provide rare insights into how variance is represented and processed in the brain, a key limitation common to them is that they only employed two discrete levels of variance (high vs. low), preventing a conceptually critical question: whether the neural representation of variability is linearly or quadratic related to evidence variability is presently unknown. It is helpful to systematically manipulate sensory evidence across several levels while keeping the average constant to answer this question. Indeed, it is possible that different neural substrates are found for both modes of representation of variability. Understanding this distinction could be crucial for refining models of decision-making and for developing a more comprehensive framework that accounts for how the brain integrates not only the mean but also the uncertainty associated with sensory evidence.

Another study that examined the neural representation of higher-order statistics in sensory evidence is the work on human brain responses to leptokurtic noise by Orbán et al. (2016) (Orbán et al., 2016). Using fMRI, they investigated how the shape of the evidence distribution— specifically, kurtosis—affects sensory processing in the human brain. Their findings showed that neural activity in the early visual cortex and higher-order decision-making regions was sensitive to the shape of the evidence distribution, with leptokurtic (high-peak, heavy-tailed) noise leading to distinct patterns of neural encoding compared to Gaussian noise. This suggests that the brain not only tracks mean but higher order summary statistics such as variance and kurtosis, potentially influencing how uncertainty is weighted in perceptual decision-making.

Here, we aim to investigate the neural correlates of variance in sensory evidence while keeping the mean constant. To do so, we revisit a previous study (Navajas et al., 2017), which examined human behaviour in making perceptual decisions under uncertainty and how individual differences shape confidence judgments. In this study, participants were presented with a sequence of 30 tilted gratings in rapid serial visual presentation and were asked to decide whether the mean orientation of the sequence was clockwise or counterclockwise relative to vertical. Crucially, while the mean orientation remained constant across all trials, the variance was systematically manipulated, allowing to isolate the effects of sensory uncertainty on behavior. Participants reported their choice and confidence in their decisions, which were used to compare several models of how uncertainty influenced metacognitive judgments. The key findings were that while most individuals’ confidence ratings reflected their estimated probability of being correct, approximately half of the participants also incorporated perceived uncertainty - irrespective of what choice they had made - in their confidence reports. These individual differences were stable over time, with some participants relying more on the probability of being correct and others integrating uncertainty more strongly into their confidence judgments. Moreover, this pattern persisted across different tasks, suggesting that individuals exhibit trait-like differences in the computational processes underlying metacognitive evaluations.

In the original study, EEG data were also collected but were not, before this work, analyzed or reported. Here, for the first time, we report the EEG findings that tell us how neural activity tracks variance in sensory evidence independent of mean. The experimental design allowed us to explore the pure electrophysiological correlates of variability and test whether EEG signals exhibit signatures of sequential accumulation of uncertainty, extending previous findings that primarily focused on confidence judgments and behavioral responses.

By analyzing this previously unreported EEG dataset, we aim to determine whether neural correlates of variance encoding are consistent with inter-individual variations in behavior, including accuracy, and metacognitive judgments. Given that previous research (Navajas et al., 2017) has shown stable individual differences in how people incorporate sensory uncertainty into their confidence reports, our analysis will test whether these behavioral tendencies are reflected in distinct neural signatures of variance assessment. Furthermore, we seek to explore whether variance-related neural activity aligns with known computational models of evidence accumulation, potentially shedding light on the neural basis of uncertainty processing in perceptual decision-making.

To promote transparency and facilitate further research, we make this dataset—including behavioral data, computational modeling code, and EEG recordings—publicly available through this paper. We hope this initiative will foster new insights into the neural and computational mechanisms underlying decision-making under uncertainty.

## Methods

We used the data obtained by Navajas et al (2017) (Navajas et al., 2017) where readers can find a more detailed description of the behavioural methodology and computational modelling. For brevity, we repeat only the key elements of the previously published methods that are directly relevant here and put more focus on detailed description of the EEG methodology. Where necessary, we have included further details in the supplementary material.

### Participants

Experiment 1 in Navajas et al. (2017) was aimed at studying the interindividual variability choice confidence perceptual decisions. We collected behavioural and EEG data from 15 participants (mean±sd age 26±8, 15 right-handed, 9 female). Participants were recruited through an advertisement at University College London and provided written informed consent. Data were collected in sessions lasting about 90 minutes each. Participants were compensated with money. All experimental procedures received approval from the research ethics committee at University College London Institute of Cognitive Neuroscience.

### Experimental protocol

#### Display, stimuli, task and procedure

Stimuli were created using the Cogent Toolbox (http://www.vislab.ucl.ac.uk/cogent.php) for MATLAB (MathWorks Inc.). Participants viewed the display on a 21-inch LCD monitor (refresh rate: 60 Hz; resolution: 1,024×768 pixels) from a distance of approximately 60 cm. In each trial, the participant viewed a series of 30 tilted Gabor patches displayed on a neutral grey background (Gaussian envelope standard deviation: 0.63°, spatial frequency: 1.57 cycles/degree, contrast: 25%). The patches were shown in rapid succession at the center of the screen, each for 200 ms, with a 50 ms interval between stimuli, resulting in an update rate of 4 Hz. At the end of the sequence, participants judged whether the average orientation of the patches was tilted clockwise or counterclockwise in relation to the vertical axis. The response options appeared as two tilted lines, one in each visual field (size: 2.2°, positioned 11.3° to the left or right of the center of the screen). The positioning of these options was randomized and counterbalanced across trials. To select the left option, participants used the ‘Q’ key on a QWERTY keyboard with their left hand, while the ‘P’ key was used to select the right option. After making their choice, participants rated their confidence in the decision on a scale from 1 to 6. A horizontal line, measuring 18.9° in length, was displayed at the center of the screen, with six evenly spaced marks indicating various confidence levels. Participants adjusted the cursor along the scale by pressing the ‘Q’ or ‘P’ keys to move it left or right, respectively. The starting point of the scale was randomly determined for each trial. After selecting a confidence rating, participants pressed the spacebar to proceed. Following a brief inter-trial interval, randomly varying between 0.7 and 0.9 seconds, a new trial began.

The orientations of the patches were drawn from uniform distributions with a fixed mean of m and endpoints at m±v. Two different means were used (m = +3 or −3 degrees) along with four different variances, determined by their endpoints: v = 10, 14, 24, or 45 degrees. These uniform distributions were sampled in a pseudorandom manner, ensuring that the mean was exactly ±3 degrees on every trial. This resulted in weak correlations, though multi-collinearity analysis showed that the presentations could not be predicted based on previous samples (R² < 0.07). The orientations were shuffled randomly to determine the order of presentation. The experiment consisted of 400 trials, with 50 trials for each of the 8 distribution types. Feedback was provided every 20 trials, displaying the number of correct trials within that block. Each block included five trials from each variance condition, presented in random order. As a result, participants could not rely on feedback to learn the performance for different variance conditions.

#### Analysis of behavioral data

To evaluate participants’ metacognitive sensitivity, we utilized the type II Receiver Operating Characteristic (ROC) curve (Hautus et al., 2021; Song et al., 2011). While alternative measures like meta-d′ are sometimes preferred, we chose the parameter-free type II AROC because it imposes fewer assumptions about the generative process underlying confidence (Fleming & Lau, 2014) and is well-established in the literature (Baird et al., 2013; Fleming et al., 2010; Song et al., 2011). Following signal detection theory, we used the area under the ROC curve (AROC) as an objective measure of metacognitive sensitivity. Confidence was indirectly assessed using a 6-point wager scale (Seth, 2008). For each wager level *i*, we calculated the probabilities *p*(*i*∣*correct*) and *p(i*∣*incorrect)*, transformed them into cumulative probabilities, and plotted them against each other with anchors at *[0,0]* and *[1,1]* (Kornbrot, 2006; Song et al., 2011). The area under the curve was computed using the method outlined by Fleming and Lau (2014) (Fleming & Lau, 2014), which accounts for Type I confounds. All analyses were conducted using Python (version 3.10.6).

#### Computational Modelling

We followed the approach presented in Navajas et al (2017) (Navajas et al., 2017) to computationally model the interindividual differences in metacognition based on three constituents: confidence in choice (i., perceived probability that a correct choice has been made), confidence in evidence (i.e., observer’s uncertainty in their estimate of the sequence mean) and baseline trait confidence (i.e., average confidence across all trials). This allows relating each participant’s psychometric profile to their corresponding neurometric profile obtained from the EEG data analysis. Below we briefly redescribe the key components of the modelling framework but refer the reader to Navajas et al (2017) (Navajas et al., 2017) for details.

Navajas et al (2017) (Navajas et al., 2017) assumed that participants track the running mean orientation of the sequence of the stimuli, updating after each new stimulus by combining a noisy estimate of the current sample with their previous mean estimate:

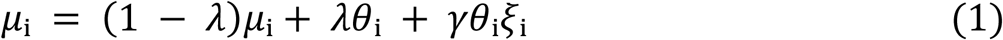

where *μ_i_* represents the estimate of the mean after *i* samples (with *μ_0_* = 0). The parameter *λ*, where *0 < λ < 1*, controls the relative weight given to recent versus earlier samples. *θ_i_* denotes the actual orientation of the *i*th, current sample in the sequence; *ξ_i_* is drawn from a standard normal distribution, and *γ* is a free parameter that specifies the perception noise. The noise’s multiplicative nature ensures that the uncertainty in updating the estimate increases with the size of the observed sample, *θ_i_*. At the end of the sequence, the participant’s choice is determined by the sign of the final mean value (*μ_30_*): if *μ_30_* is positive, they choose clockwise, and if μ_30_ is negative, they choose anticlockwise.

Navajas et al. (2017) (Navajas et al., 2017) proposed that participants also compute the variance of the sensory evidence, such that the perceived variance, denoted *σ_30_²*, is defined by:

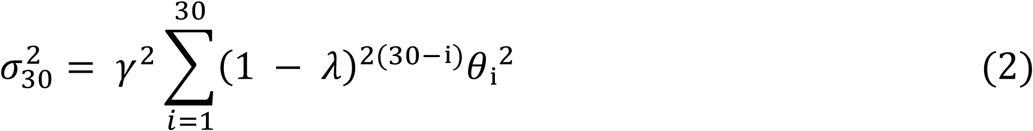

For any agent that employs this stochastic updating model (equations 1 and 2) confidence in choice is readily calculated as the shaded area under the Gaussian distribution in Fig. 1c which corresponds to the perceived probability of being correct (see equation 5-9 below). Similarly, confidence in evidence is given by the Fisher information (i.e., 1/*σ_30_²*) on each trial.

**Figure 1.**
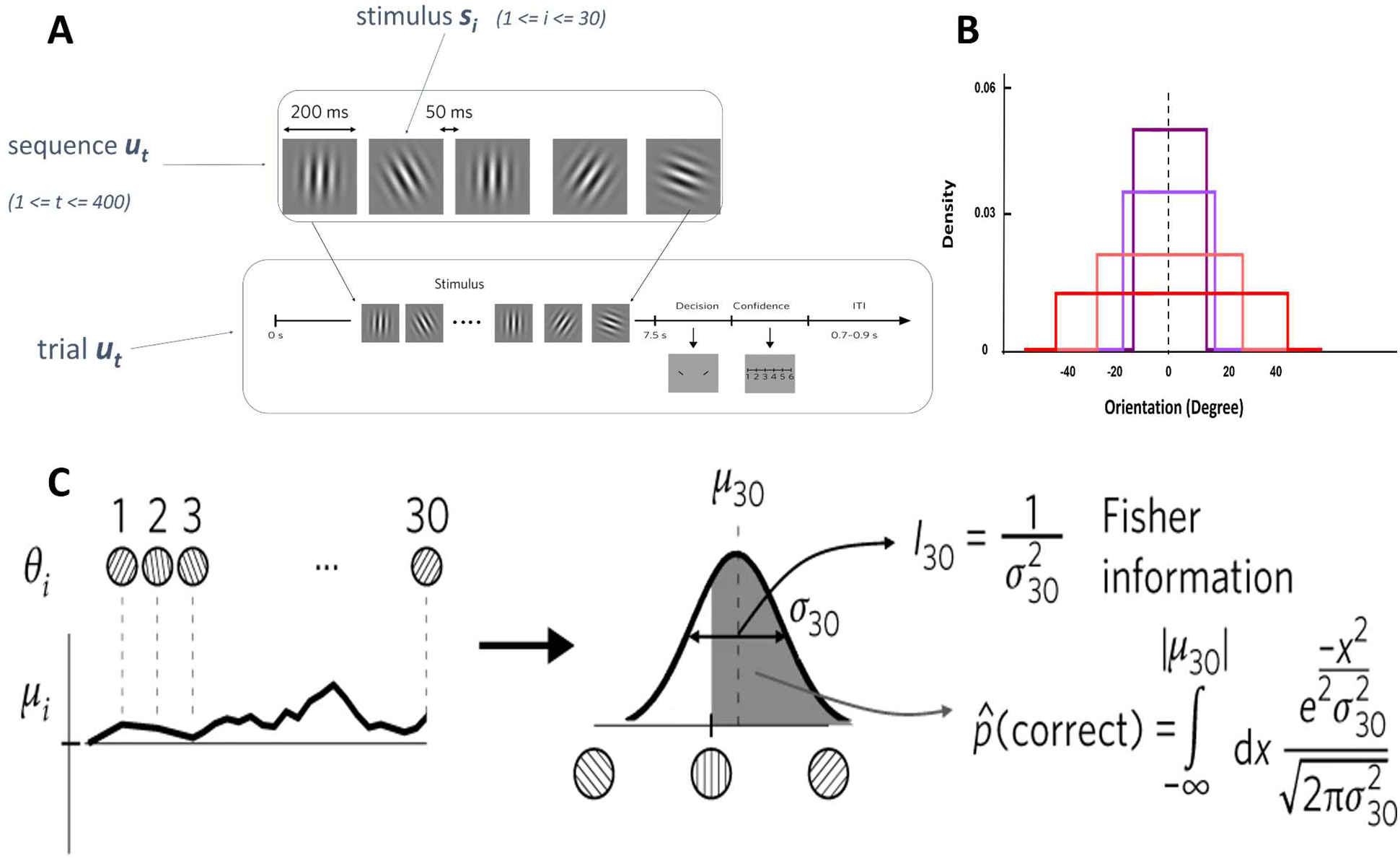
Experimental paradigm and confidence estimating model. (A) In each trial, participants viewed a rapid stream of 30 tilted Gabor patches, flashed at the fovea and updated at 4 Hz. Their task was to judge whether the average orientation across the sequence was tilted to the left or right, followed by a confidence rating. Trials were separated by an inter-trial interval (ITI) randomly drawn between 0.7 and 0.9 seconds. (B) The orientations were sampled from a uniform distribution with a fixed mean of either +3° or −3°. The variability of the stimulus was manipulated by changing the range (*m* ± *v*), where v was set to 10°, 14°, 24°, or 45°, resulting in four distinct variance conditions. (C) On each frame, the current estimate (*μᵢ*) was updated by integrating the previous estimate with a noisy observation of the current Gabor (*θᵢ*). The black line illustrates one instance of this running integration. After the 30th sample, a binary choice was made based on the sign of *μ₃₀*. Subjective confidence (i.e., the probability of being correct) and the Fisher information were then derived.

#### Estimating the Fisher information and the perceived probability of being correct

Following Navajas et al (2017), using the best-fitting parameters *λ* and *γ* obtained from the stochastic updating model (the values that maximize *L(λ, γ)* in equation (2)), we estimated, for each trial, both the observed Fisher information and the expected perceived probability of making a correct decision. The observed Fisher information is simply the inverse variance of the participants’ estimate, which is computed using equation (2) (Fig. 1.c.). The expected perceived probability of making a correct decision, denoted as d, is given by:

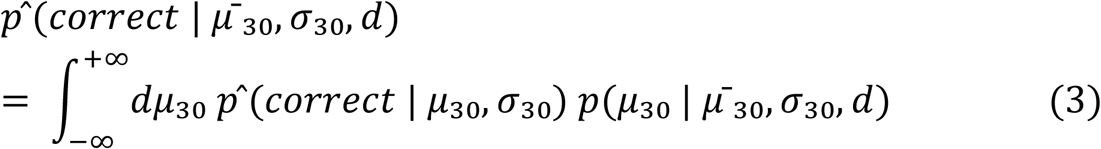

The first term inside the integral, *P (correct | μ_30_, σ_30_)*, represents the shaded area under the Gaussian curve in Fig. 1c. Therefore, it is described by the cumulative normal distribution,

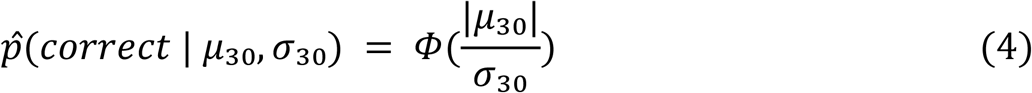

The second term in the integral, *p(μ_30_ | μ ^-^₃₀, σ_30_, d)*, reflects the likelihood of observing μ_30_given μ^-^ ₃₀, σ_30_, and the decision, d. If the decision is clockwise (*d* = +1), μ_30_is constrained to be positive, while for an anticlockwise decision (*d* = −1), μ_30_must be negative. These conditions are captured using the Heaviside step function, *Θ(x)*, which takes the value 1 when *x* > 0 and 0 otherwise, leading to:

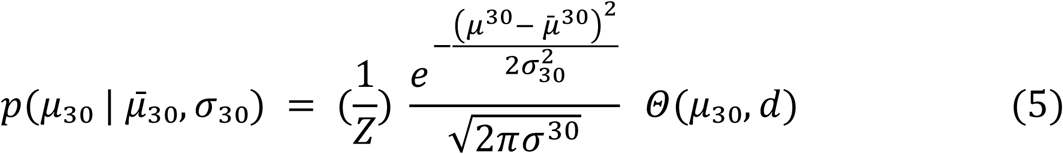

where *Z* is the normalization constant,

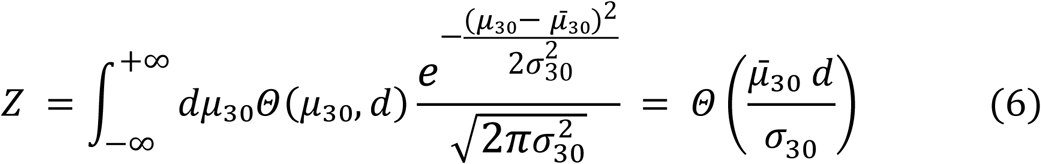

By combining these two expressions, we obtain:

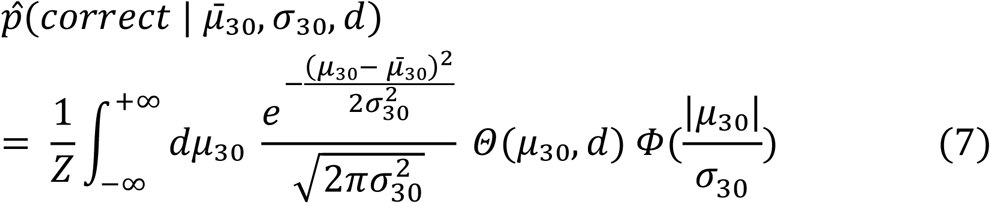

The expected perceived probability of being correct (equation (7)) depends on the decision, d, whereas the Fisher information (equation (2); Fig. 1c) is independent of d and thus choice-independent. On each trial, *p(μ_30_ | μ ^-^₃₀, σ_30_, d)* was computed numerically using Matlab.

#### Deconstructing confidence into its metacognitive components

For each individual, we performed a multivariate ordinal regression (McCullagh, 1980). This regression fits a logistic model with fixed effects and different offsets for each of the five possible splits in the rating scale.

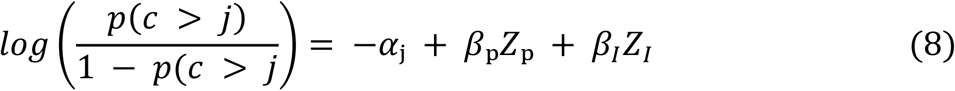

Where 1 ≤ *j* ≤ 5, c represents confidence, and *Z_p_* and *Z_I_* are the z-scored estimates of the perceived probability of being correct and Fisher information on each trial. The outputs of this regression include the offsets *α_1_*, …, *α_5_*, as well as the weights *β_p_* and *β_I_*. To summarize the computations involved in confidence, we selected α_3_ (the offset when the scale is split into halves, referred to as the overall confidence), *β_p_* (the weight of the probability of being correct on confidence), and *β_I_* (the weight of information on confidence).

### EEG

#### EEG acquisition and preprocessing

Signals were recorded using a BIOSEMI Active Two 64-channel EEG system adhering to the 10– 20 system of electrode placement on the scalp. The sampling rate was set to 256 Hz and channels were referenced to the right-mastoid electrode

The raw data were processed with EEGLAB software (Delorme & Makeig, 2004). First, a notch filter was applied between 45 and 55 Hz to eliminate line noise. To remove high-frequency interference, an FIR filter covering 0.1 - 100 Hz was used. Artifacts were detected and removed through visual inspection, supplemented by independent component analysis. Noisy trials were discarded through visual checks, and any channels affected by noise were interpolated using EEGLAB. The signals were segmented into distinct epochs, each aligned with the stimulus presentation, spanning from 2000 ms before the stimulus onset to 2500 ms after stimulus offset, for a total duration of 12,000 ms (with 7,500 ms allocated to the stimulus presentation). Following preprocessing, epochs of EEG data that exceeded (or were below) a threshold of 100 (–100) μV were excluded from the analysis (for detailed data analysis, refer to Table 6 and Materials and Methods)(Kelly & O’Connell, 2013). For regional analysis of the EEG data, the recorded channels were grouped into five brain regions based on anatomical and functional considerations. These regions and their corresponding channels are as follows:

##### Frontal Region

This includes the channels AF7, AF3, AF4, AF8, AFz, F1, F3, F5, F7, F2, F4, F6, F8, and Fz, which predominantly capture activity from the frontal brain areas.

##### Lateral-Parietal Region

Channels P1, P3, P5, P7, P9, P2, P4, P6, P8, and P10 are grouped in this region, covering the lateral parietal cortex.

##### Centro-Parietal Region

Comprising the channels CP1, CP2, CP3, CP4, CPz, and Pz, this region reflects activity from the central and parietal areas.

##### Occipital Region

The occipital channels O1, O2, Oz, POz, PO7, PO3, PO4, and PO8 are linked to the visual cortex and surrounding areas.

##### Temporal Region

This region includes the channels FT7, FT8, T7, T8, TP7, and TP8, corresponding to activity in the temporal lobes. This regional segmentation allowed for localized analysis of brain activity, enhancing the interpretation of spatially specific neural dynamics.

We analyzed EEG data during the 7,500 ms stimulus presentation and applied a 500 ms baseline correction immediately prior to stimulus onset. Given that the inter-trial interval ranged from 700 to 900 ms, a longer pre-stimulus window would have included residual activity from the previous trial. Restricting the baseline period to 500 ms helped minimize this overlap and provided a cleaner reference baseline. For each trial, we averaged the selected channels within each brain region and computed the mean signal amplitude and the mean power using Morlet wavelet analysis. We excluded a total of five subjects from the analysis: three due to poor-quality EEG data and two due to mismatches between EEG trials and behavioral data.

We explored the relationship between the stimulus standard deviation and variance and the mean values (amplitude and power) of the EEG signals using a mixed-effects model, and computed coefficients for each region. Additional details on these models are available in the supplementary materials.

## Results

### Behavior

Our behavioral analysis was consistent with the previous report (Navajas et al 20217), replicating the relationship between accuracy, confidence, and variance (See Supplementary Materials Fig. S1.A-C). To assess metacognitive sensitivity, we calculated the area under the ROC curve (AUC) for each participant (Fleming & Lau, 2014). AUC is driven from Signal Detection Theory and reflects the individual’s ability to differentiate between correct and incorrect responses by expressing their confidence flexibly. The mean AUC score across participants was 0.635 ± 0.058 (Fig. 2), indicating considerable variation among participants.

**Figure 2.**
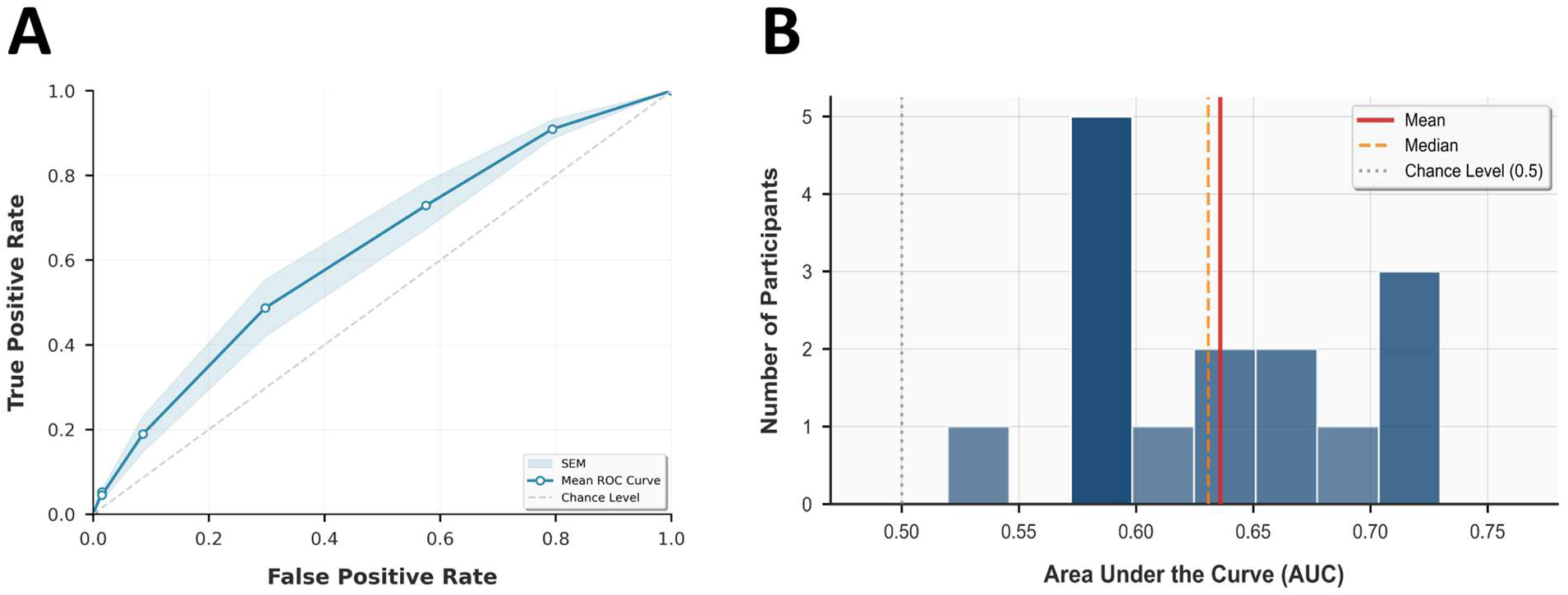
**(A)** Metacognitive sensitivity (AUC) is calculated by constructing the type 2 Receiver Operating Characteristic (ROC) function, where the false alarm rate is on the x-axis and the hit rate on the y-axis for each given confidence criterion. Type 2 decisions are judgments about one’s own accuracy, such as confidence ratings or deciding whether a previous decision was correct. The area under the curve indexes metacognitive sensitivity. (B) Empirical distribution of AUC among our cohort of participants.

### EEG

#### EEG Signal Power and Frequency Peaks

We began our EEG signal analysis by plotting the power spectrum (Fig. 3A), which revealed three prominent peaks. The first peak, occurring around 4 Hz, corresponds directly to the 250 ms sensory update cycle used in the experiment (200 ms stimulus followed by a 50 ms inter-stimulus interval), yielding four updates per second.

**Figure 3.**
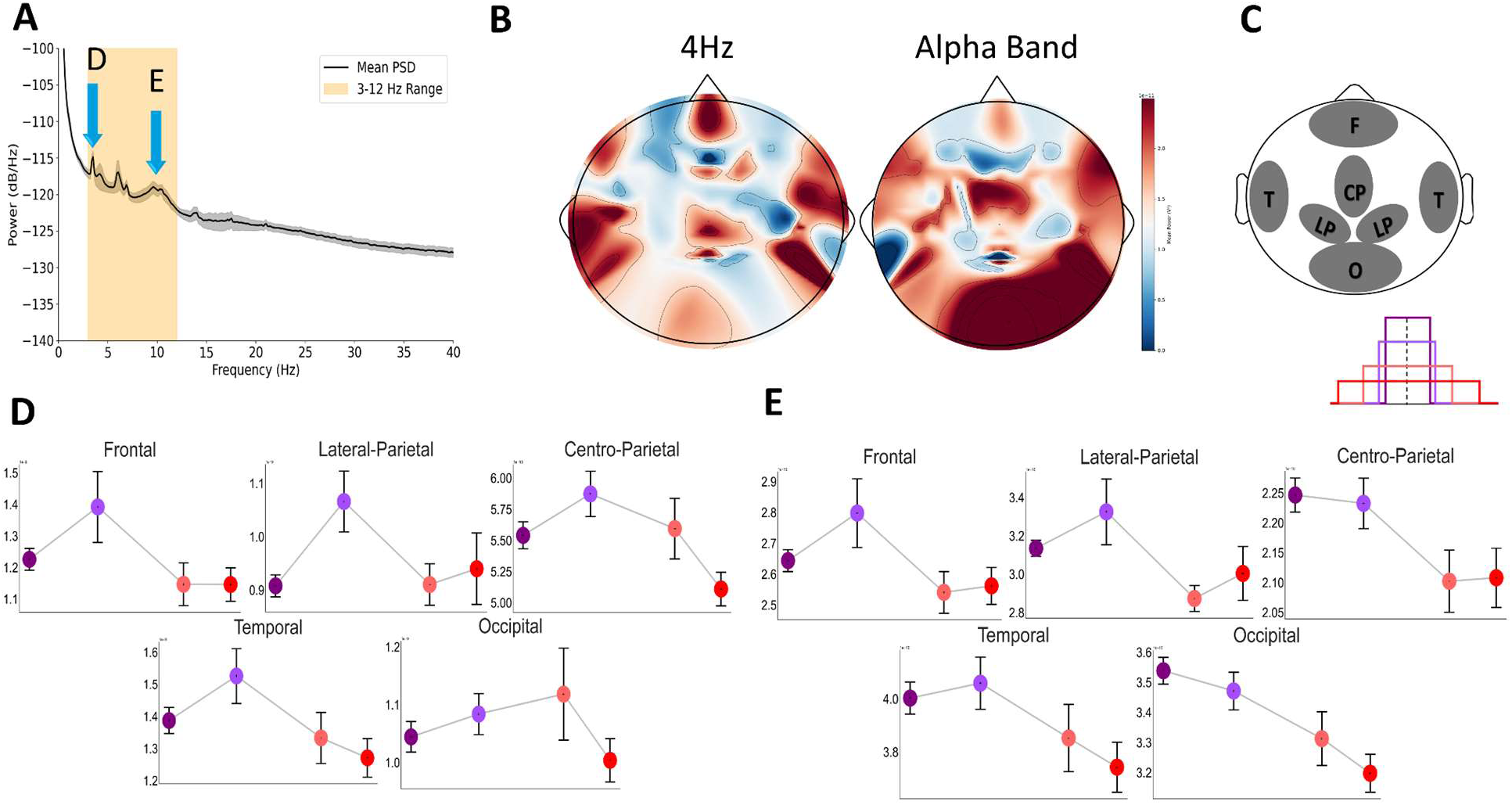
Neural correlates of variance of sensory stimulus. A. Power Frequency Histogram. B. Topographical Map of Power Across Brain Regions for a Specific Frequency (4 Hz). C. Selected Channels for five interested Regions. The colors (purple, lavender, salmon, red) indicate a range from low to high variance. D. Averaged mean powers across varying levels of variance are shown for five brain regions. E. the same as D panel but for alpha band (9-11 Hz).

**Figure 4.**
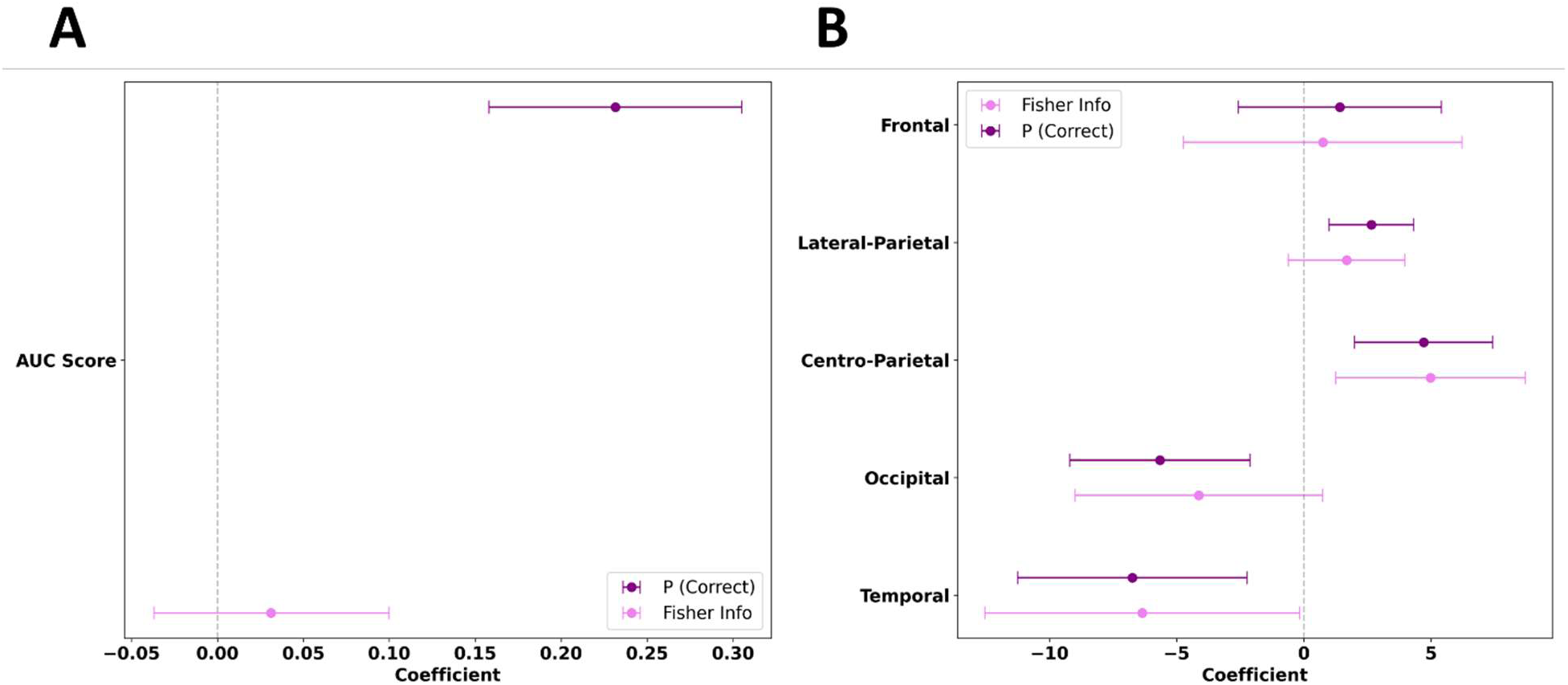
Computational and neural correlates of interindividual differences in metacognitive accuracy (AUC). Exploring Neural Correlates of Variance and Model Parameters as Predictors of Metacognitive Sensitivity. A. The model’s parameters are used as predictors for the AUC score. B. Relationship between beta coefficient of P(correct) and the Fisher Information associated with the beta coefficients within each region.

#### Sensitivity to Sensory Variability

To examine how neural responses tracked variability in the stimulus sequence, we compared the average EEG signal power across four levels of standard deviation in the sensory input. We observed a clear quadratic relationship between signal power and stimulus variability (*β_Var_Frontal_* = −1.73e-12, *P* = 0.008, *β_Var_Lateral_Parietal_* = −7.36e-13, *P* = 0.051, *P* = 0.008, *β_Var_Centro_Parietal_* = −4.79e-13, *P* < 0.001, *β_Var_Occipital_* = −6.51e-13, *P* = 0.061, *β_Var_Temporal_* = −1.29e-12, *P* = 0.020), particularly in Frontal, Centro-Pareital and Temporal regions where effects were significant (Fig. 2D and supplementary figure S3). This quadratic pattern suggests that the brain’s processing of sensory evidence is most sensitive to moderate levels of variance, while responses to very high variance are less pronounced. Importantly, this trend does not co-vary with task difficulty or attentional demand, and therefore cannot be explained by factors such as error monitoring, cognitive effort, or the strength of the sensory signal. In the regions showing significant effects (frontal, centro-parietal, and temporal), this pattern of response may reflect the brain’s adaptive mechanisms for efficiently allocating resources to stimuli that are neither too predictable nor too unpredictable.

#### Theta Activity and Task Engagement

The second peak in the spectrum fell within the theta band (approximately 5.5–7.5 Hz), a range typically linked to cognitive functions such as visual processing, integration of information, and decision-making (Cavanagh & Frank, 2014; Herweg et al., 2020; Jacobs et al., 2006). This aligns well with the demands of our task, which required averaging Gabor orientations, and echoes findings from previous studies (Kawasaki et al., 2014; Su et al., 2024). Moreover, the periodic 4Hz presentation of stimuli may have contributed to enhanced theta synchronization in this range (Clouter et al., 2017; Tomassini et al., 2017). In line with earlier findings, we also observed a quadratic relationship between theta power and task parameters (*β_Var_Frontal_* = −4.85e-13, *P* = 0.014, *β_Var_Lateral_Parietal_* = −2.80e-13, *P* = 0.054, *P* = 0.008, *β_Var_Centro_Parietal_* = −1.30e-13, *P* = 0.002, *β_Var_Occipital_* = −2.44e-13, *P* = 0.009, *β_Var_Temporal_* = −3.85e-13, *P* = 0.020; Figure S6 and Table S4).

#### Alpha Power and Task Difficulty

The third peak was observed in the alpha band (9–11 Hz), which is well-established as being sensitive to task difficulty, attentional demand, and cognitive load (Jensen & Mazaheri, 2010; Klimesch, 1999). In our data, increasing stimulus variability—interpreted here as increased task difficulty—was associated with a linear decrease in alpha power in the centro-parietal, Temporal and occipital regions (*β_Std_Frontal_* = −4.557e-13, *P* = 0.2404, *β_Std_Lateral_Parietal_* = −1.002e-12, *P* = 0.096, *P* = 0.008, *β_Std_Centro_Parietal_* = −8.414e-13, *P* < 0.001, *β_Std_Occipital_* = −1.645e-12, *P* < 0.001, *β_Std_Temporal_* = −1.353e-12, *P* < 0.001; Fig. 2E and Table S5). This contrasts with the quadratic pattern seen in the lower frequency bands (Figure 2D), suggesting distinct underlying mechanisms for how these frequency bands respond to cognitive demands.

### Inter-individual differences

#### Computational correlates of interindividual differences in metacognitive sensitivity

To identify which computational features support accurate confidence monitoring, we used the stochastic updating model (Navajas et al., 2017), which estimates a computational phenotype for each participant—indicating whether their confidence reports rely more on subjective probability of being correct or on Fisher information. We then regressed individual AUC scores (a measure of metacognitive sensitivity; Fig. 2) on these model-derived weights. The analysis revealed that only the weight for subjective probability (*βₚ*) significantly predicted AUC (*β_1_* = 0.2465 ± 0.05, *P* < 0.001; Fig. 3A, Table S1), while the weight for Fisher information (*β_FI_*) did not (*β_2_* = 0.064 ± 0.084, *P* = 0.461; Fig. 3A, Table S1). This suggests that individual differences in metacognitive sensitivity are best explained by the degree to which confidence tracks subjective probability rather than raw sensory precision.

#### Neural correlates of interindividual differences in metacognitive sensitivity

Next, we examined whether inter-individual variability in model parameters—subjective probability (*βₚ*) and Fisher information (*β_FI_*)—could be predicted by neural responses to sensory variance across brain regions. Specifically, we analyzed whether mean 4 Hz power modulated by stimulus variance could account for differences in these computational weights.

We found that neural correlates of *β_FI_* were significant only in the centro-parietal region (Fig. 3B and Table S5), making it the key neural predictor of Fisher information use in confidence estimation. In contrast, βₚ showed significant neural associations in all regions except the frontal cortex (Fig. 3B and Table S6). When each region was tested separately (Tables S7–S14), centro-parietal and occipital regions showed opposite trends: Fisher information correlated positively with neural power, while P(correct) correlated negatively. These results suggest distinct neural signatures for the two sources of confidence and highlight the central role of the centro-parietal region in supporting metacognitive computations.

## Discussion

The central aim of this study was to examine the neural signature of sensory evidence variability in the human brain under two important conditions. First, throughout the experimental conditions, the mean of the sensory evidence was held constant. Second, we examined variability at four different levels to permit the observation of both linear and nonlinear neural coding of variability. These two issues are important because previous research has extensively examined how the mean strength of sensory evidence influences decision-making. Although a few studies have indeed explored how the brain encodes the variability—or uncertainty—of sensory inputs (Bang & Fleming, 2018; Orbán et al., 2016; Zylberberg et al., 2016), these studies have invariably focused on 2 levels of variability i.e. low vs high which, by design, does not permit distinguishing between linear and nonlinear trends. Our experiment was particularly geared to address these two issues.

Our key findings are two folds. First, we found that evidence for neural coding of sensory variability was observed in the EEG signal power but not in the event related potentials (see Supplementary material). Second, we found that these neural correlates demonstrated considerable systematic heterogeneity across different frequency bandwidths and different scalp locations.

Given the nature of our stimulus (i.e., 4Hz rapid serial visual presentation of high contrast gratings), we observed strong EEG entrainment at this frequency and its higher harmonics. We found that the power of these stimulus-specific entrainments bore a quadratic relationship to sensory variability most strongly pronounced in the frontal and centro-parietal regions. Importantly, the quadratic pattern indicates that the neural response was not simply driven by the range of the stimulus distribution but was instead tuned to variance as a distinct statistical property. This sensitivity to variance, rather than just the spread of sensory inputs, suggests a refined encoding of sensory uncertainty.

Meanwhile, a different form of covariation was observed between variability and EEG power in the EEG alpha band, particularly in the centro-parietal and occipital regions. Here, alpha power in these areas were linearly related to variability with a systematic decrease in alpha power in higher variability conditions. These latter findings align with previous research linking alpha suppression to increased task difficulty, more cognitive effort and increased uncertainty. Pilipenko and Samaha (2024) (Pilipenko & Samaha, 2024) showed that reductions in alpha power track the accumulation of sensory evidence, with lower alpha power reflecting greater engagement in decision-making processes. Their findings suggest that alpha suppression reflects a state of heightened sensitivity to incoming evidence, facilitating accumulation dynamics. Similarly, Chen et al. (2023) (Chen et al., 2023) found that alpha oscillations play a key role in integrating sensory uncertainty, with alpha suppression enhancing sensitivity to variable sensory input. Our findings are consistent with these findings and thus help push the field one step forward in better characterizing the role of alpha power in the brain’s handling of cognitive load. However, none of these studies can establish whether alpha suppression plays a causal role in enhancing sensitivity under conditions of increased difficulty or whether it is merely a downstream consequence of cognitive processes that regulate uncertainty, serving as an epiphenomenal byproduct rather than an active mechanism. Addressing this distinction would require causal interventions, such as transcranial alternating current stimulation (tACS) to manipulate alpha oscillations directly, or neurofeedback approaches to determine whether voluntary modulation of alpha affects perceptual sensitivity.

### Neural and computational Interindividual differences in metacognition

Previous studies that investigated the neural correlates of interindividual differences in metacognitive sensitivity (Baird et al., 2013; Fleming et al., 2010) correlations between variations in metacognition and measures of neural connectivity (Baird et al., 2013) or structure (Fleming et al., 2010) that did not relate, directly or indirectly, to representation and processing of variability and uncertainty by the human brain. It seems logical that one would have started the search for these correlates by examining the neural processes underlying variability and uncertainty. Towards this goal, here we reported our participants’ metacognitive sensitivity using the AUC measure, permitting the readers to directly compare the results with previous studies. Moreover, we then asked if such sensitivity was in any meaningful way represented in the neural correlates of variability identified here. However, in order to better disentangle the computational and cognitive constructs contributing to the role variability in behavior and brain, we made use of our own earlier computational framework (Navajas et al., 2017).

The framework proposed by Navajas et al. (2017) (Navajas et al., 2017) offered a principled method to dissect the individuals’ confidence judgments into distinct computation of variability: (1) the precision of the sensory evidence, captured by Fisher information, which is independent of the choice made, and (2) the subjective probability of being correct, which is inherently tied to the decision. By modeling these related but conceptually separable components, the framework allowed for subject-specific inference of how much each factor contributed to metacognitive sensitivity. This approach enabled a richer understanding of confidence as a personalized construct rather than a uniform readout, and it provides the foundation for our current investigation into how neural correlates of variability might support these distinct computational contributions.

We demonstrate that the conventional metric of metacognitive sensitivity (AUC) loads primarily on the subjective probability component and not on Fisher Information. This permitted combining our model-based interpretation of metacognition behaviour with the neural correlates of variability with one another and the behaviour and the EEG signal could be related to one another through their link to variability of sensory evidence. We found some modest but significant correlations between the model-based participant-specific parameters —subjective probability (*βₚ*) and Fisher information (*β_FI_*) – and variance-related neural activity, most clearly observed in the centro-parietal region. Importantly, the CPP relationship to variability was found to be correlated with both model components even though only one of them i.e., subjective probability (*βₚ*) was connected to metacognitive sensitivity. These findings suggest that interindividual differences in metacognition are not only reflected in brain structure (Fleming et al., 2010) and connectivity (Baird et al., 2013) but also in dynamic neural encoding of uncertainty during decision-making. Future research could integrate these approaches to examine whether the structural and resting-state correlates identified in earlier studies predict variance-related EEG activity during perceptual tasks.

## Acknowledgments

BB, GL and JE were supported by the European Research Council (ERC) under the European Union’s Horizon 2020 research and innovation programme (819040 - acronym: rid-O). BB is also supported by the EMERGE EIC (Project 101070918).

## Supplementary Materials

### Extended Materials and Methods and Results

We analyzed how the regression weights for p(correct) and Fisher information impacted the AUC score by applying the following regression model (Eq. S1):

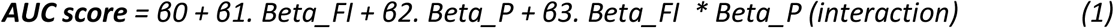

Where *Beta_P* is the regression weights for p(correct). *Beta_FI* refers to the regression weights forFisher information.

**Table S1.** Details of statistical results in the relationship between AUC and beta_P and betaFi.

| Regressors | Estimate | SE | CI<br>[0.025<br>0.975] | t-Stat | p-Value | No.<br>Observations |
| --- | --- | --- | --- | --- | --- | --- |
| **Const**<br>**(β<sub>0</sub>)** | 0.5118 | 0.024 | [0.459 0.564] | 21.429 | <0.001 | 15 |
| **Beta_FI**<br>**(β<sub>1</sub>)** | 0.064 | 0.084 | [-0.121<br>0.249] | 0.763 | 0.461 | 15 |
| **Beta_P**<br>**(β<sub>2</sub>)** | 0.2465 | 0.05 | [0.137 0.356] | 4.962 | <0.001 | 15 |
| **Interaction**<br>**(β<sub>3</sub>)** | -0.0577 | 0.136 | [-0.356<br>0.241] | -0.426 | 0.679 | 15 |

### Behavioral data

Consistent with previous studies, our behavioral analysis revealed that participants performed better as variance decreased (Fig. S1.A). Additionally, confidence on correct trials increased significantly as variance dropped (Fig. S1.B). However, on error trials, confidence remained relatively unaffected by variance (Fig. S1.B). Our data did not show the pattern that had been predicted based on normative arguments, at least not on average. As illustrated in Fig. XB, while confidence on correct trials did increase as variance decreased, on error trials, confidence remained relatively unaffected by variance. Our analysis of reaction time did not reveal a significant effect of variance changes (Fig. S1.C).

As outlined, we employed the area under the ROC curve (AUC) as an objective measure of metacognitive sensitivity. The AUC score quantifies an individual’s ability to accurately differentiate between correct and incorrect responses based on their confidence ratings. The mean AUC score across all subjects was 0.635 ± 0.058 (Fig. S1.D), indicating a moderate level of metacognitive sensitivity. The standard deviation (±0.058) suggests variability in participants’ metacognitive accuracy, with some individuals demonstrating better self-awareness in assessing the correctness of their responses, while others show less confidence or greater inaccuracy in their self-assessments. This variability reflects the differences in metacognitive ability across participants, implying that some individuals are more adept at evaluating their decisions than others.

It is important to note that the AUC score is far from 1.0 (which would indicate perfect discrimination between correct and incorrect responses), suggesting considerable room for improvement in participants’ metacognitive judgments. These finding highlights that, on average, participants were overconfident or misjudging the correctness of their responses, particularly in error trials. For instance, they should have been less confident on low-variance error trials compared to high-variance error trials, as their likelihood of being correct was lower in the former. This discrepancy suggests that, overall, participants struggled to calibrate their confidence in relation to the actual probability of correctness, especially when variance in the evidence was low.

**Figure S1.**
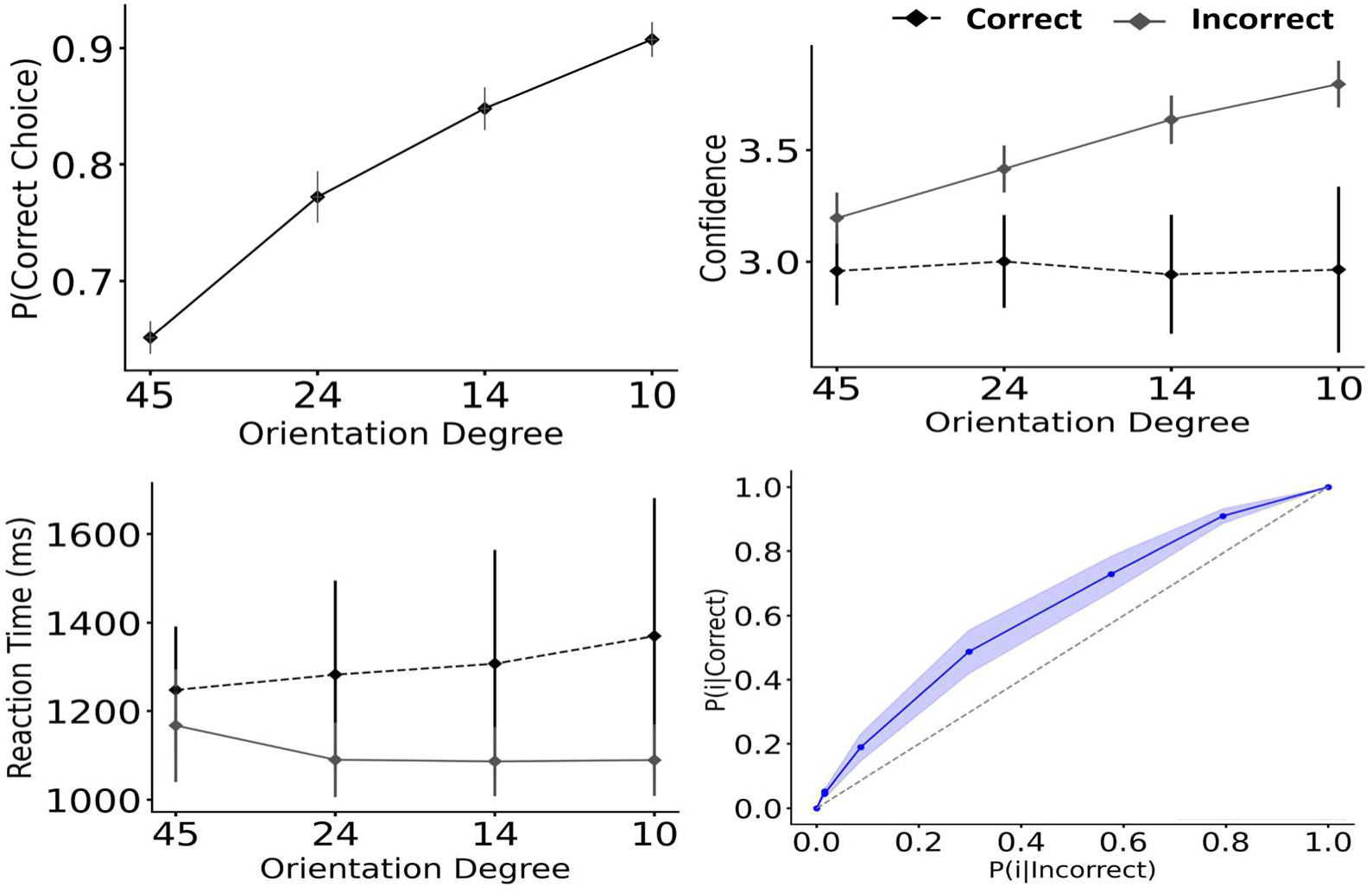
Behavioral results. A. Accuracy. B. Confidence. C. Reaction time. D. Metacognitive sensitivity

The model was fitted individually for each participant, and this fitting process did not influence participant exclusion based on poor EEG data or misalignments. To ensure our dataset remains comparable to Navajas (2017), we demonstrate that these 15 participants exhibit behavior indistinguishable from the 15 participants who did not undergo EEG recording (P β_P = 160, P β_FI = 160).

**Figure S2.**
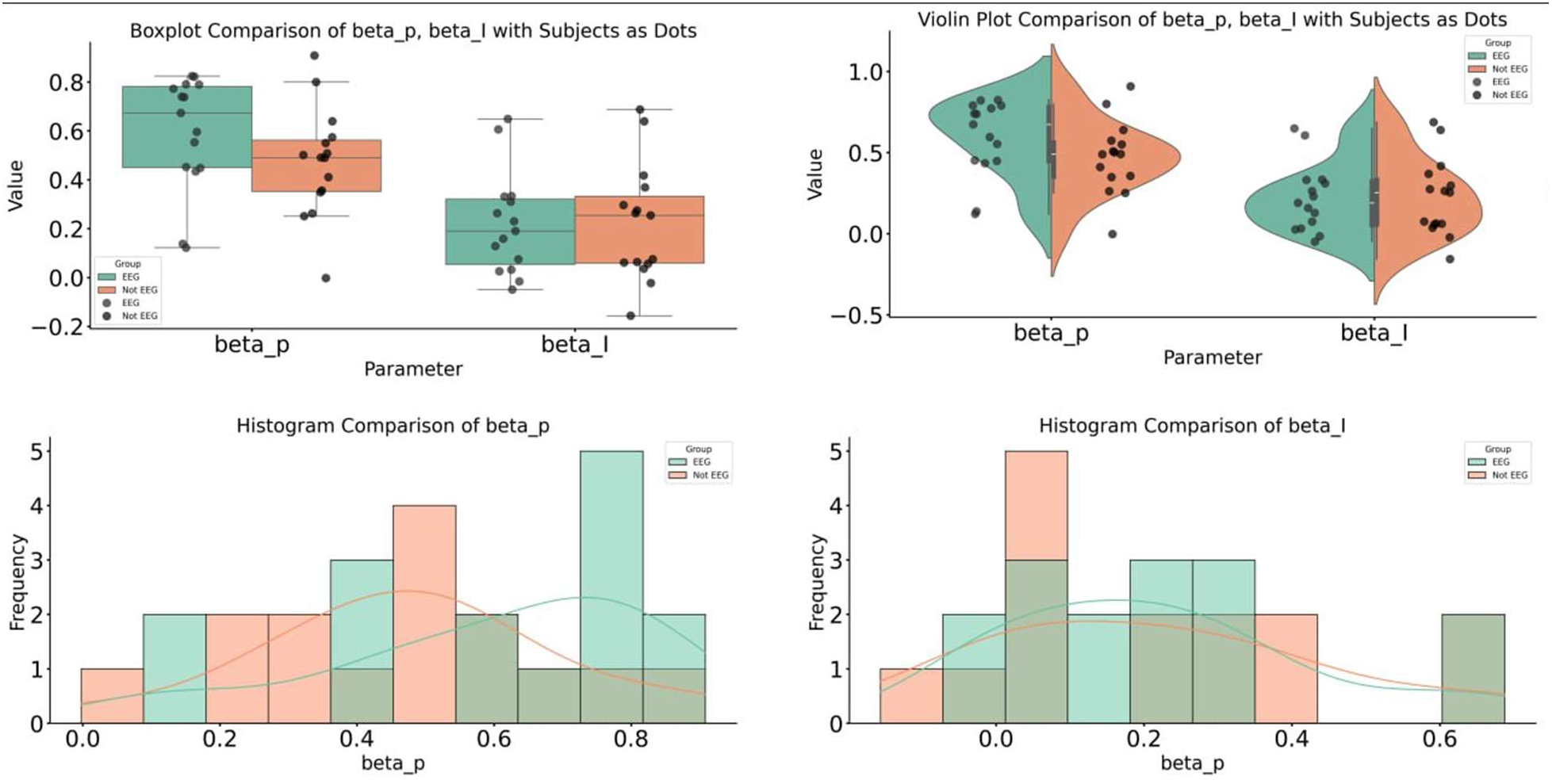
Comparing the distributions of beta_p and beta_I for the 15 subjects we analyze and the 15 we don’t. Up Left: Boxplot. Up Right: Violinplot. Bottom Left: Histogram of Beta_P. Bottom Right: Histogram of Beta_I.

**Figure S3.**
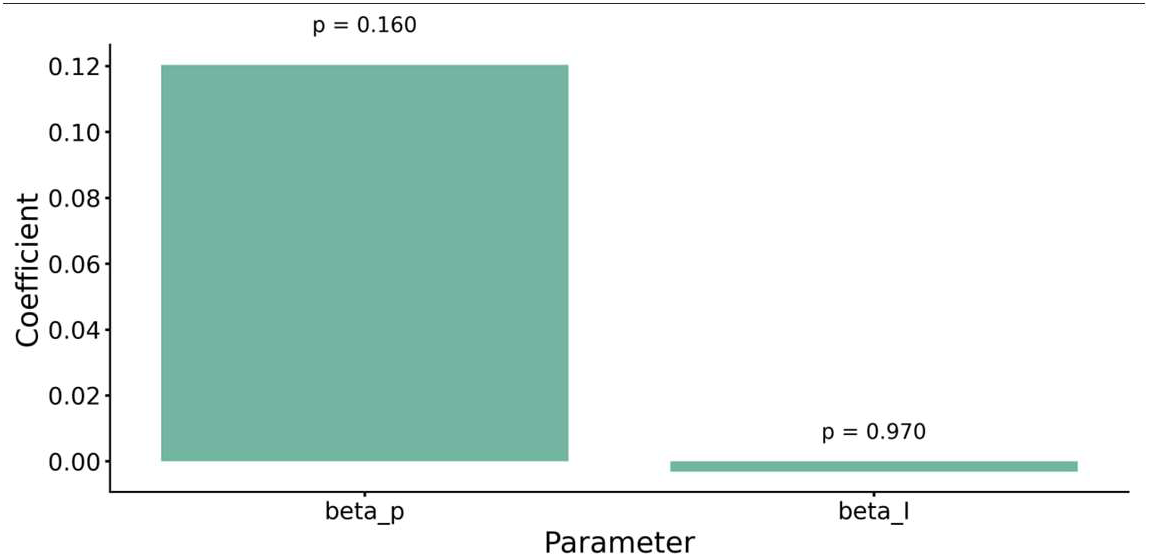
Statistical test of EEG vs Non-EEG, Regression Coefficients for beta_p and beta_I along with p-values.

### Neural correlate of mean amplitude

As previously outlined, a duration of 7,500 ms was allocated for the stimulus presentation. For each subject, we calculated the average signal across all regions (Fig. S4.A). Next, we averaged the signals over time for each level of low to high variance. However, the results did not reveal any specific pattern (Fig. S4.B).

**Figure S4.**
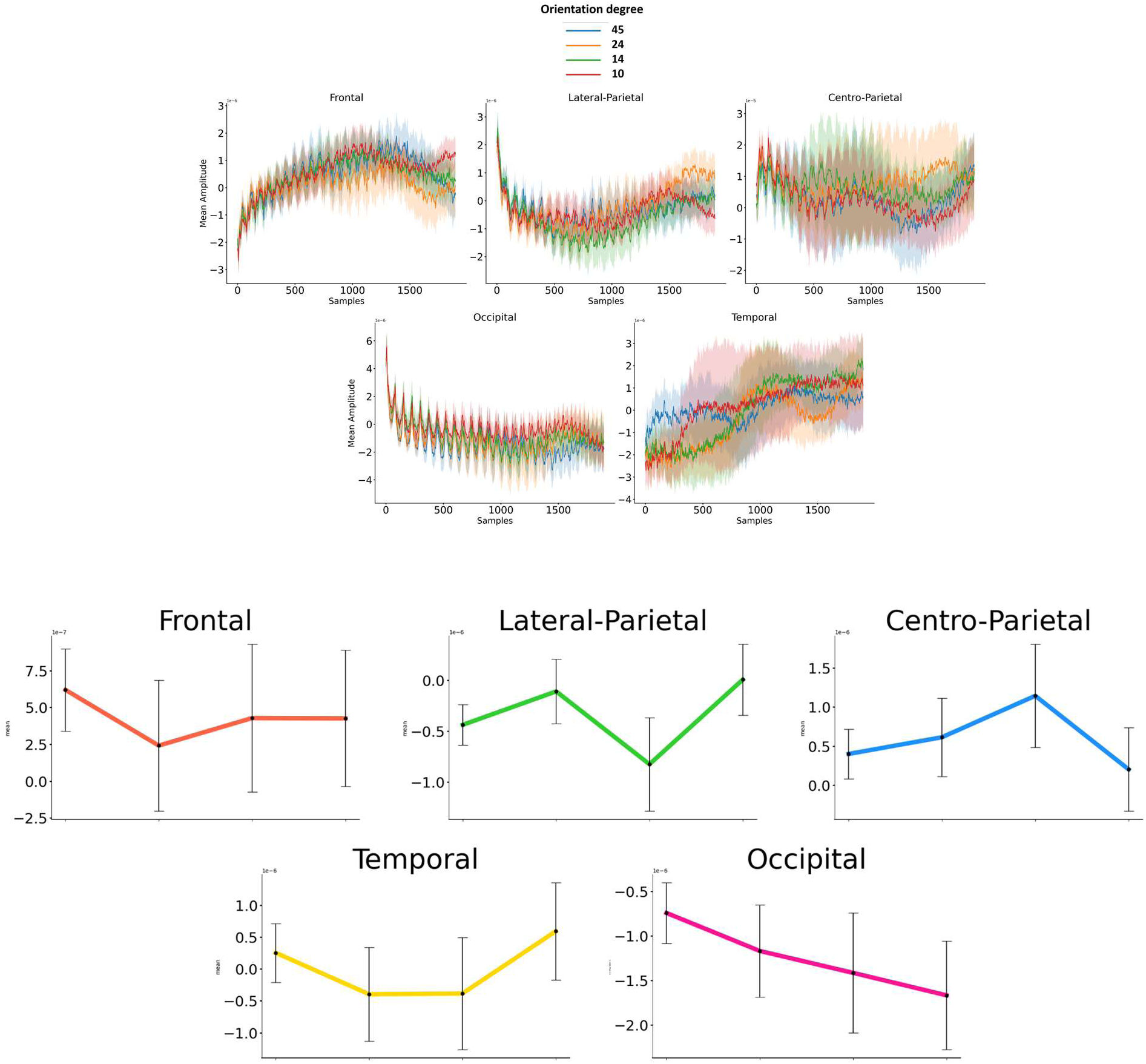
Top: Mean amplitude of preprocessed signals across orientation levels for each region from stimulus onset to offset. Bottom: Neural correlates of variance of sensory stimulus for Mean amplitude. Error bars are standard errors of the mean (SEM) across subjects.

### Neural correlate of Power Frequency

We investigated the connection between the stimulus standard deviation and variance and the mean values (amplitude and power) of the EEG signals using a mixed-effects model (Eq. S2):

**Figure S5.**
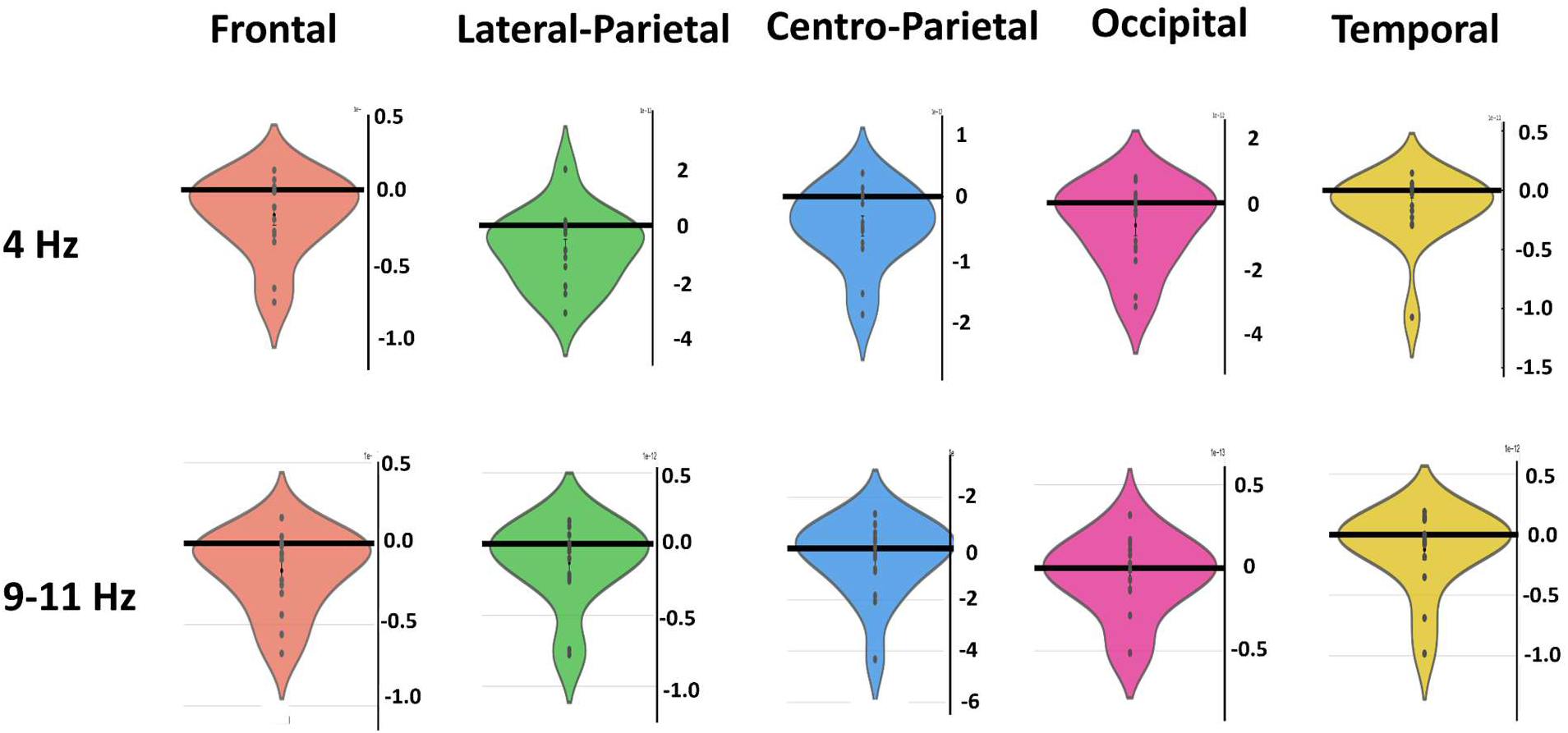
Beta Coefficient for the Quadratic Term: The violin plots on the right side of each panel illustrate the distribution of the beta coefficient for the quadratic term for five brain regions (red: Frontal, yellow: Temporal, pink: Occipital, green: Lateral-Parietal, and blue: Centro-Parietal), with individual dots representing subjects

**Figure S6.**
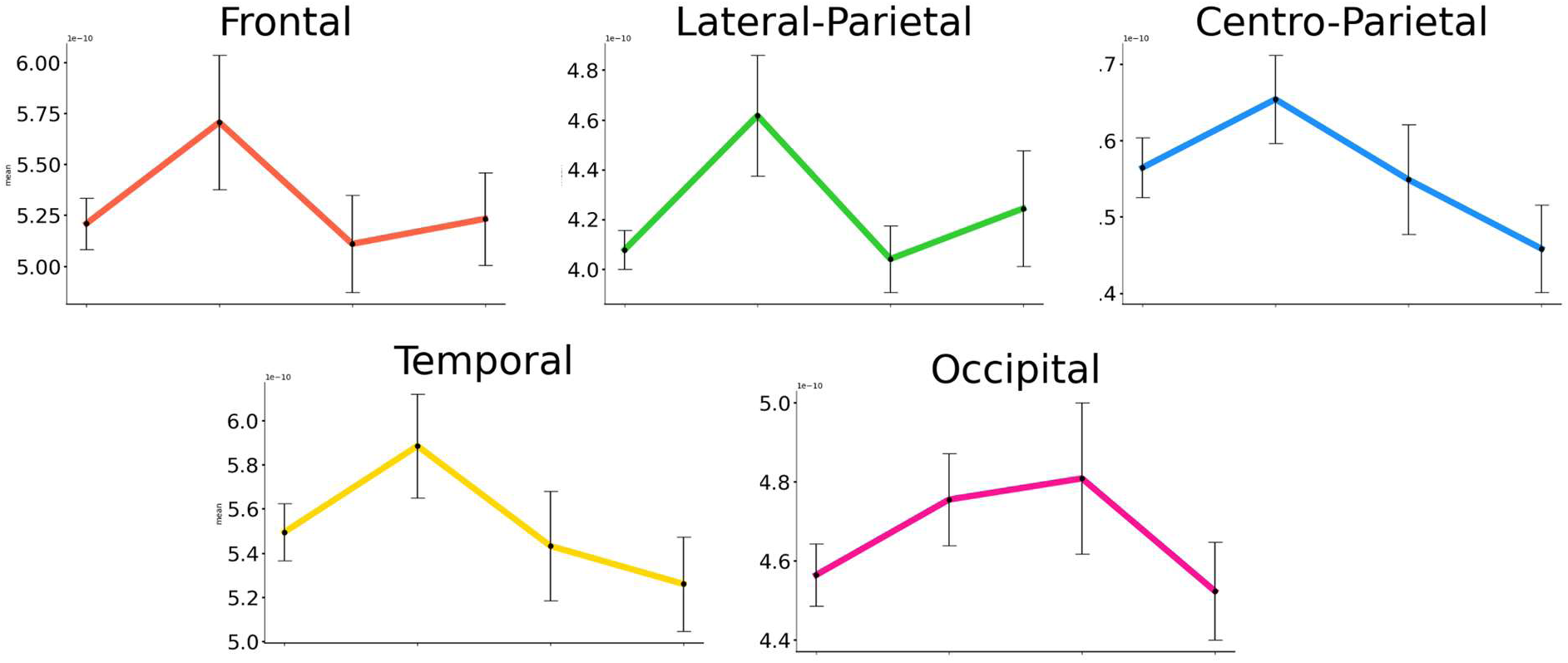
Neural correlates of variance of sensory stimulus in the second bump (5.5-7.5 HZ). Averaged mean powers across varying levels of variance are shown for five brain regions (red: Frontal, yellow: Temporal, pink: Occipital, green: Lateral-Parietal, and blue: Centro-Parietal).

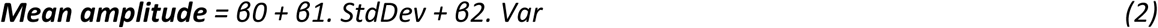

Where *StdDev* is the standard deviation of stimulus sets and *Var* refers to the variance of stimulus sets.

The same mixed-effects model (Eq. S2) was applied to the mean power of the signal at 4 Hz.

**Table S2.** Details of statistical results in the relationship between regions and coefficients of mean of signal amplitude.

| Regions | Regressors | Coefficients | P-value |
| --- | --- | --- | --- |
| **Frontal** | **StdDev**<br>**($\beta_1$)** | **-5.48e-08** | **0.681** |
|  | **Var**<br>**($\beta_2$)** | **1.56e-09** | **0.696** |
| **Lateral-Parietal** | **StdDev**<br>**($\beta_1$)** | **-6.63e-08** | **0.497** |
|  | **Var**<br>**($\beta_2$)** | **2.04e-9** | **0.485** |
| **Centro-Parietal** | **StdDev**<br>**($\beta_1$)** | **1.77e-07** | **0.244** |
|  | **Var**<br>**($\beta_2$)** | **-5.22e-09** | **0.252** |
| **Occipital** | **StdDev**<br>**($\beta_1$)** | **-7.96e-08** | **0.624** |
|  | **Var**<br>**($\beta_2$)** | **7.86e-10** | **0.872** |
| **Temporal** | **StdDev**<br>**($\beta_1$)** | **-3.68e-07** | **0.096** |
|  | **Var**<br>**($\beta_2$)** | **1.13e-08** | **0.087** |

**Figure S7.**
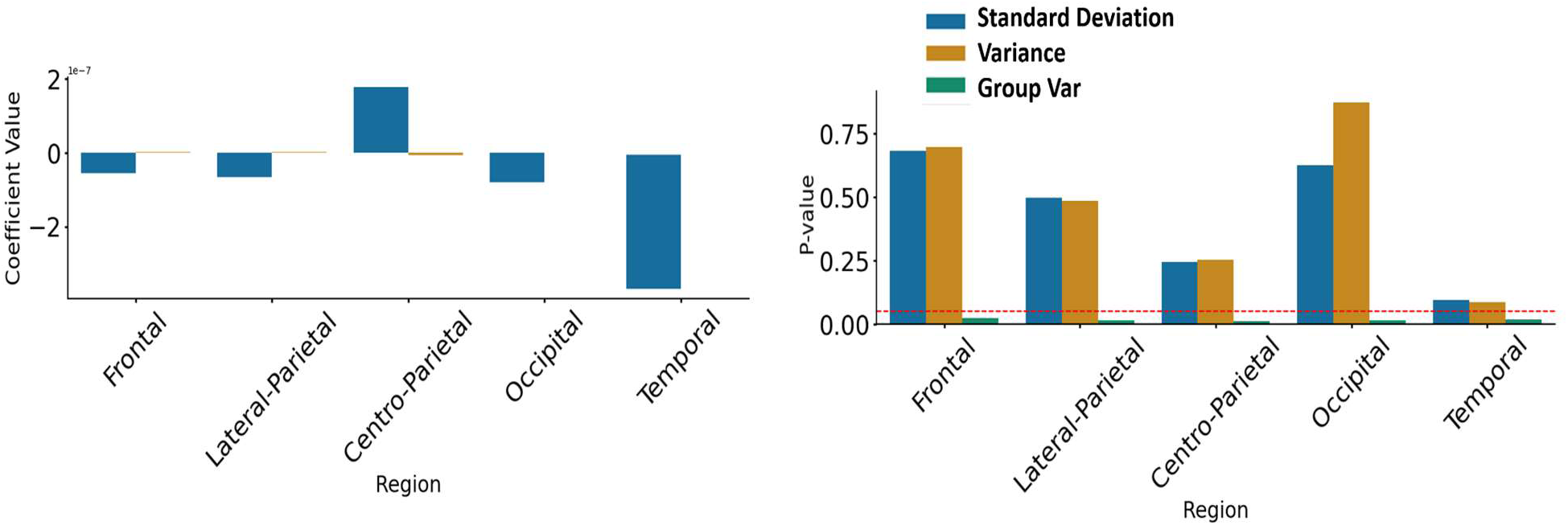
Details of statistical results for mean amplitude. A. P-value. B. Coefficients

**Figure S8.**
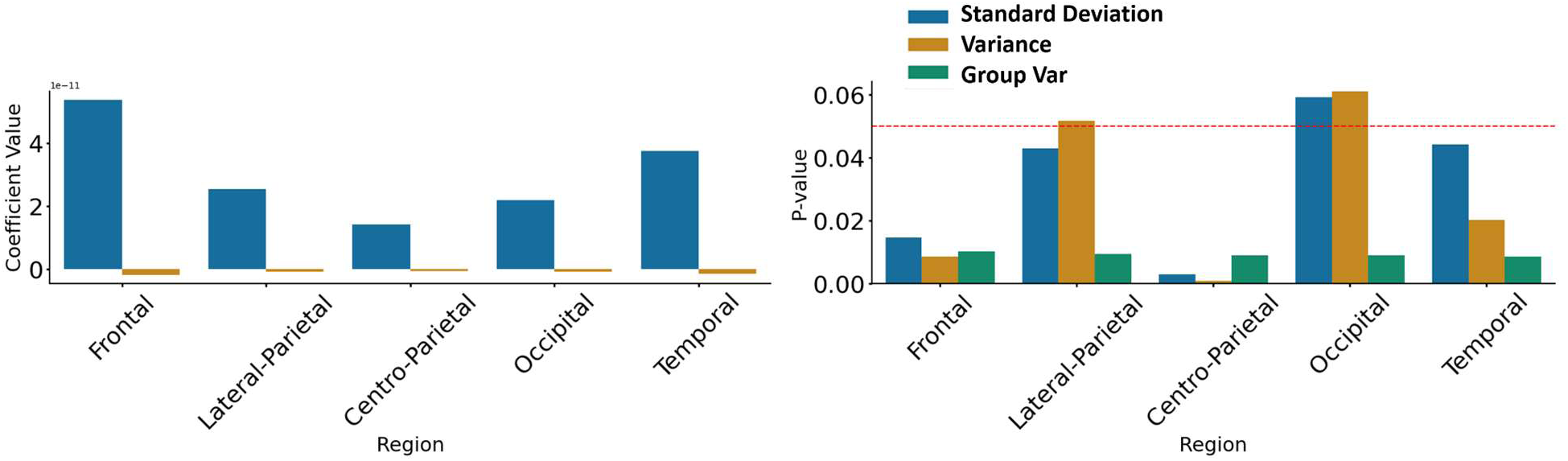
Details of statistical results for mean power. A. P-value. B. Coefficients

**Table S3.** Details of statistical results in the relationship between regions and coefficients of mean of signal power (4 HZ).

| Regions | Regressors | Coefficients | P-value |
| --- | --- | --- | --- |
| **Frontal** | **StdDev**<br>**($\beta_1$)** | **5.37e-11** | **0.014** |
|  | **Var**<br>**($\beta_2$)** | **-1.73e-12** | **0.008** |
| **Lateral-Parietal** | **StdDev**<br>**($\beta_1$)** | **2.55e-11** | **0.042** |
|  | **Var**<br>**($\beta_2$)** | **-7.36e-13** | **0.051** |
| **Centro-Parietal** | **StdDev**<br>**($\beta_1$)** | **1.42e-11** | **0.02** |
|  | **Var**<br>**($\beta_2$)** | **-4.79e-13** | **<0.001** |
| **Occipital** | **StdDev**<br>**($\beta_1$)** | **2.18e-11** | **0.059** |
|  | **Var**<br>**($\beta_2$)** | **-6.51e-13** | **0.061** |
| **Temporal** | **StdDev**<br>**($\beta_1$)** | **3.75e-11** | **0.044** |
|  | **Var**<br>**($\beta_2$)** | **-1.29e-12** | **0.020** |

### Neural correlate of mean power (5.5 to 7.5 Hz)

Neural correlates of variance of sensory stimulus in the second bump (5.5-7.5 HZ).

**Table S4.**
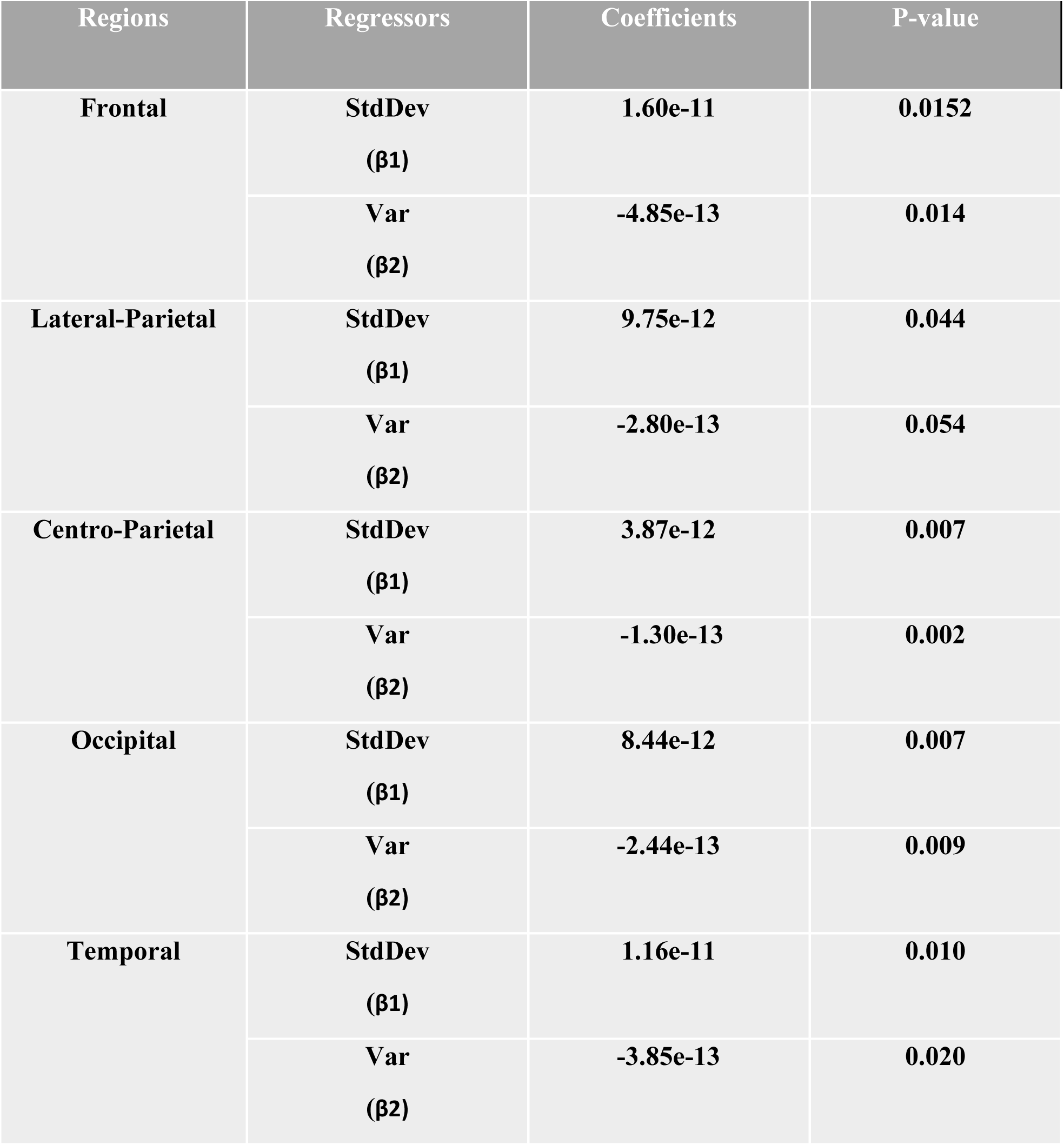
Details of statistical results in the relationship between regions and coefficients of mean of signal power (5.5 - 7.5 HZ).

| Regions | Regressors | Coefficients | P-value |
| --- | --- | --- | --- |
| **Frontal** | **StdDev**<br>**($\beta_1$)** | **1.60e-11** | **0.0152** |
|  | **Var**<br>**($\beta_2$)** | **-4.85e-13** | **0.014** |
| **Lateral-Parietal** | **StdDev**<br>**($\beta_1$)** | **9.75e-12** | **0.044** |
|  | **Var**<br>**($\beta_2$)** | **-2.80e-13** | **0.054** |
| **Centro-Parietal** | **StdDev**<br>**($\beta_1$)** | **3.87e-12** | **0.007** |
|  | **Var**<br>**($\beta_2$)** | **-1.30e-13** | **0.002** |
| **Occipital** | **StdDev**<br>**($\beta_1$)** | **8.44e-12** | **0.007** |
|  | **Var**<br>**($\beta_2$)** | **-2.44e-13** | **0.009** |
| **Temporal** | **StdDev**<br>**($\beta_1$)** | **1.16e-11** | **0.010** |
|  | **Var**<br>**($\beta_2$)** | **-3.85e-13** | **0.020** |

**Figure S9.**
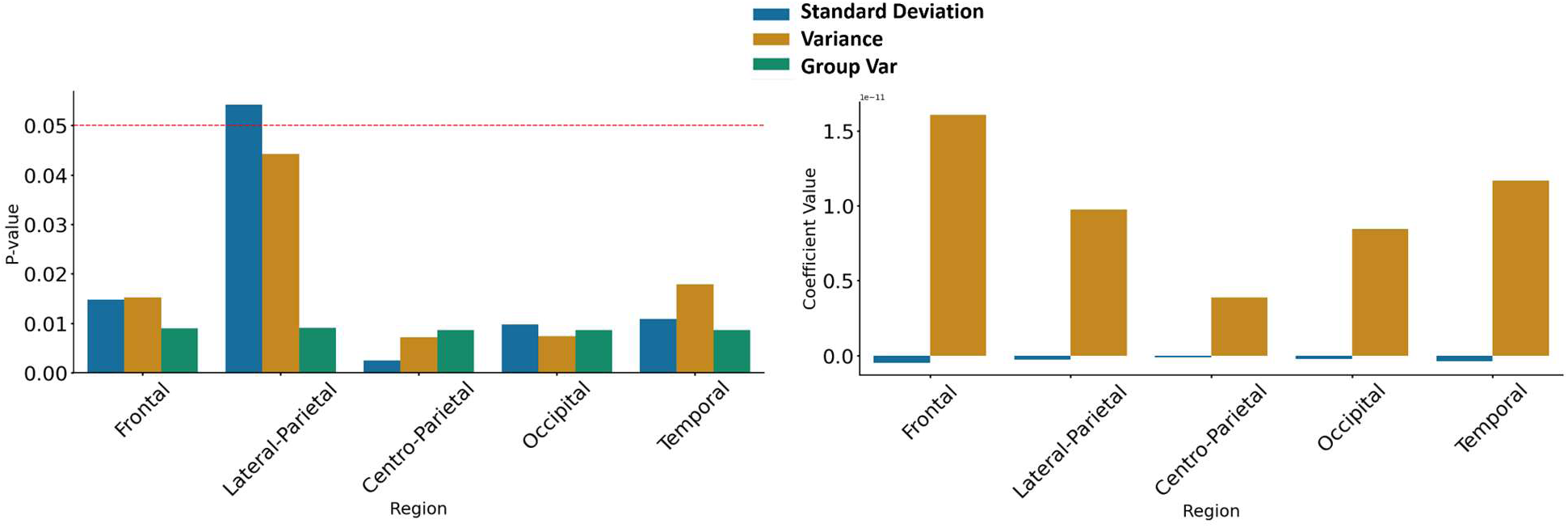
Details of statistical results for mean power in 5.5-7.5 Hz. A. P-value. B. Coefficients

### Neural correlate of mean power (9 to 11 Hz)

We expanded the frequency band range from 4 Hz to 9-11 Hz, encompassing the Alpha band, to investigate the neural correlates of variance within this range. Our results revealed that the Centro-parietal, Occipital and Temporal region showed significant association.

**Table S5.**
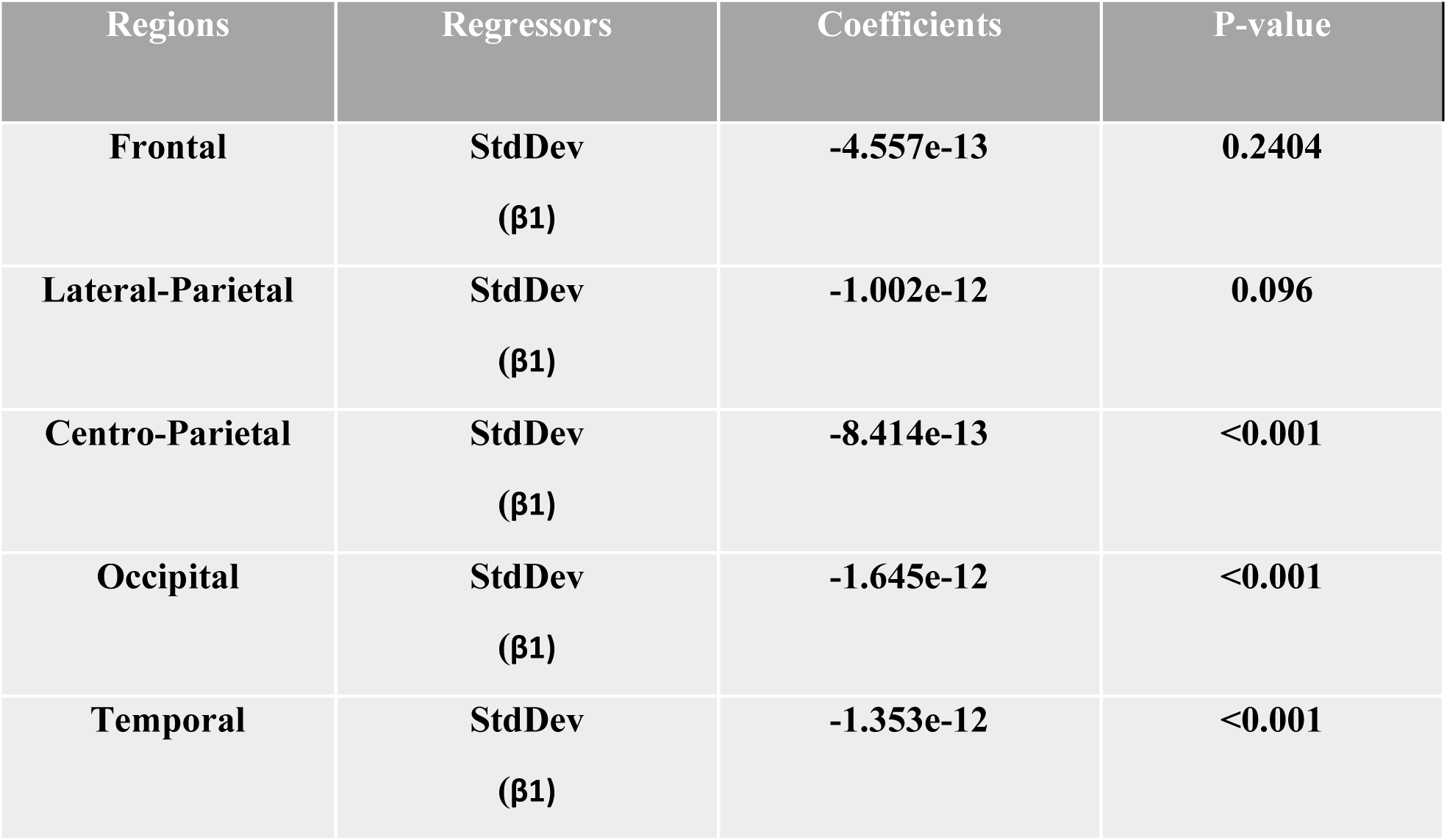
Details of statistical results in the relationship between regions and coefficients of mean of signal power (9 - 11 HZ).

**Figure S10.**
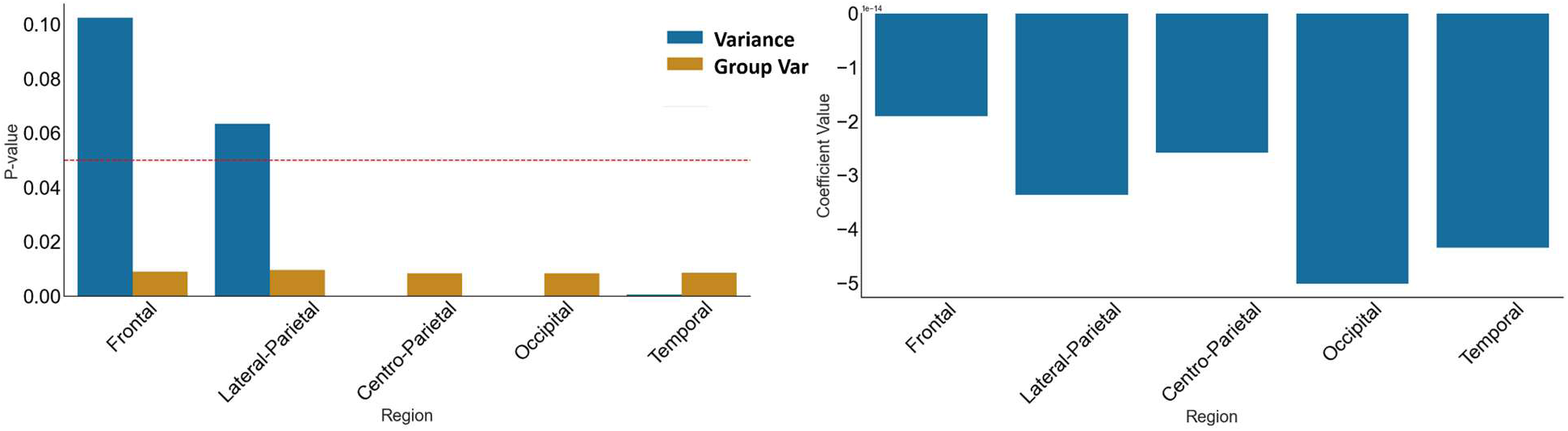
Details of statistical results for mean power in 9-11 Hz. A. P-value. B. Coefficients

### Relation between R_AUC and Beta_P and Beta_FI

We used pairwise correlation to examine the relationship between the AUC score and the model parameters (Beta_P and Beta_FI). Our results indicate a strong correlation between the AUC score and Beta_P (r = 0.91) and a moderate correlation with Beta_FI (r = 0.45).

**Figure S11.**
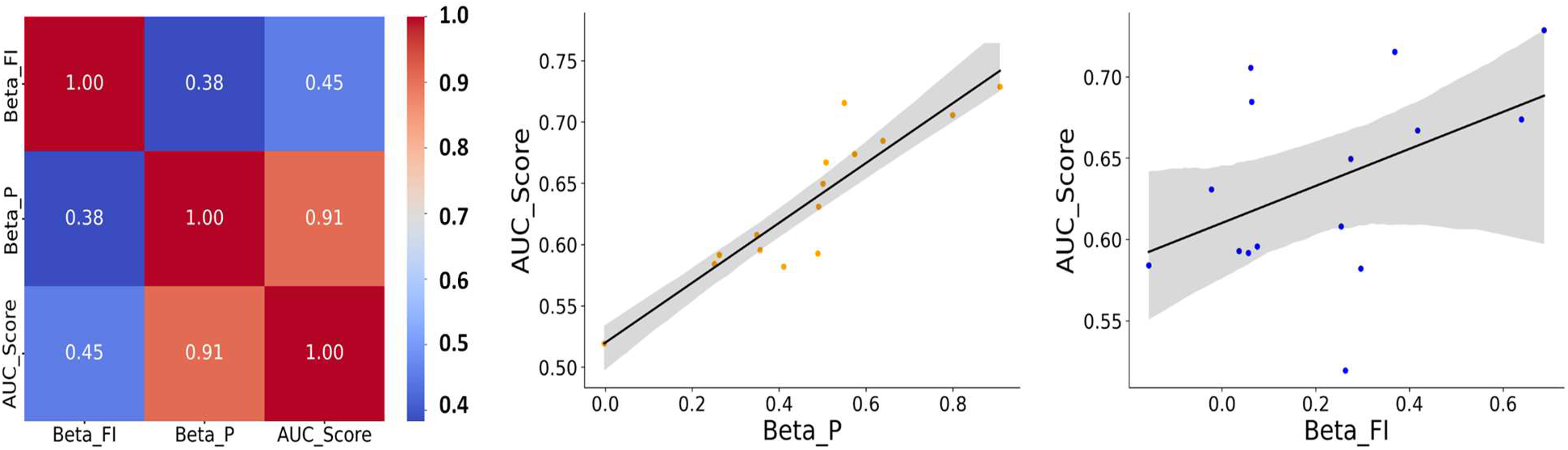
Relation between R_AUC and Beta_P and Beta_FI. A. pair-wised correlation. B. samples for each parameter

### Relation between Beta_P and Beta_FI with EEG beta of variance

We examined the influence of EEG coefficients, including the standard deviation and variance of stimuli from various brain regions, on the beta of Fisher information and the probability of a correct response (P(Correct)), using the following regression model (Eq. S3):

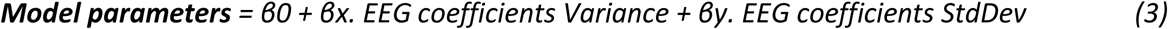

Where *x,y = (Navajas et al., 2017,5)* and x are the coefficients of different regions for variance and y are the standard deviations. Model parameters are Beta_P and Beta_FI.

**Table S6.** Details of statistical results in the relationship between fisher information and EEG beta of variance and standard deviation.

| *Regressors* | *coef* | *std err* | *t* | *P>\|t\|* | *[0.025* | *0.975]* | *No. Observations* |
| --- | --- | --- | --- | --- | --- | --- | --- |
| **Intercept** | **0.2209** | **0.041** | **5.337** | **0.006** | **0.106** | **0.336** | **15** |
| **Frontal (Var)** | **0.7478** | **1.976** | **0.378** | **0.724** | **-4.74** | **6.235** | **15** |
| **Lateral-Parietal (Var)** | **1.6837** | **0.825** | **2.041** | **0.111** | **-0.606** | **3.974** | **15** |
| **Centro-Parietal (Var)** | **4.9821** | **1.343** | **3.708** | **0.021** | **1.252** | **8.712** | **15** |
| **Occipital (Var)** | **-4.1291** | **1.753** | **-2.355** | **0.078** | **-8.996** | **0.738** | **15** |
| **Temporal (Var)** | **-6.3585** | **2.233** | **-2.848** | **0.047** | **-12.558** | **-0.159** | **15** |
| **Frontal (StdDev)** | **0.2999** | **1.997** | **0.15** | **0.888** | **-5.244** | **5.843** | **15** |
| **Lateral-Parietal (StdDev)** | **1.8814** | **0.878** | **2.142** | **0.099** | **-0.558** | **4.32** | **15** |
| **Centro-Parietal (StdDev)** | **4.4332** | **1.215** | **3.648** | **0.022** | **1.059** | **7.808** | **15** |
| **Occipital (StdDev)** | **-3.6906** | **1.609** | **-2.294** | **0.084** | **-8.158** | **0.776** | **15** |
| **Temporal (StdDev)** | **-6.112** | **2.214** | **-2.761** | **0.051** | **-12.259** | **0.035** | **15** |

**Table S7.** Details of statistical results in the relationship between beta of P(correct) and EEG beta of variance and standard deviation.

| *Regressors* | *coef* | *std err* | *t* | *P>\|t\|* | *[0.025* | *0.975]* | No.<br>Observati<br>ons |
| --- | --- | --- | --- | --- | --- | --- | --- |
| ***Intercept*** | **0.4724** | **0.03** | **15.665** | **<0.001** | **0.389** | **0.556** | **15** |
| ***Frontal (Var)*** | **1.4185** | **1.44** | **0.985** | **0.38** | **-2.58** | **5.417** | **15** |
| ***Lateral-Parietal (Var)*** | **2.6554** | **0.601** | **4.418** | **0.012** | **0.987** | **4.324** | **15** |
| ***Centro-Parietal (Var)*** | **4.7117** | **0.979** | **4.814** | **0.009** | **1.994** | **7.429** | **15** |
| ***Occipital (Var)*** | **-5.6611** | **1.277** | **-4.432** | **0.011** | **-9.207** | **-2.115** | **15** |
| ***Temporal (Var)*** | **-6.747** | **1.627** | **-4.147** | **0.014** | **-11.264** | **-2.23** | **15** |
| ***Frontal (StdDev)*** | **0.7384** | **1.455** | **0.508** | **0.638** | **-3.301** | **4.777** | **15** |
| ***Lateral-Parietal (StdDev)*** | **2.7675** | **0.64** | **4.324** | **0.012** | **0.99** | **4.545** | **15** |
| ***Centro-Parietal (StdDev)*** | **4.3211** | **0.886** | **4.88** | **0.008** | **1.862** | **6.78** | **15** |
| ***Occipital (StdDev)*** | **-5.04** | **1.172** | **-4.299** | **0.013** | **-8.295** | **-1.785** | **15** |
| ***Temporal (StdDev)*** | **-6.6708** | **1.613** | **-4.136** | **0.014** | **-11.149** | **-2.192** | **15** |

To investigate the relationship between EEG variance coefficients and the model parameters for each region separately, we applied the following linear regression:

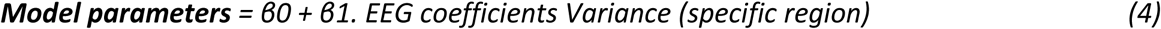

**Table S8.** Details of statistical results in the relationship between beta of P(correct) and EEG beta of variance (Centro-Parietal).

| *Regressors* | *coef* | *std err* | *t* | *P>\|t\|* | *[0.025* | *0.975]* | *No. Observations* |
| --- | --- | --- | --- | --- | --- | --- | --- |
| Intercept | 0.4724 | 0.059 | 8.064 | <.001 | 0.346 | 0.599 | 15 |
| Centro-Parietal (Var) | -0.0416 | 0.059 | -0.711 | 0.49 | -0.168 | 0.085 | 15 |

**Table S9.** Details of statistical results in the relationship between beta of P(correct) and EEG beta of variance (Lateral-Parietal).

| *Regressors* | *coef* | *std err* | *t* | *P>\|t\|* | *[0.025* | *0.975]* | *No. Observations* |
| --- | --- | --- | --- | --- | --- | --- | --- |
| Intercept | 0.4724 | 0.06 | 7.912 | <.001 | 0.343 | 0.601 | 15 |
| Lateral-Parietal (Var) | -0.0011 | 0.06 | -0.018 | 0.986 | -0.13 | 0.128 | 15 |

**Table S10.** Details of statistical results in the relationship between beta of P(correct) and EEG beta of variance (Occipital).

| *Regressors* | *coef* | *std err* | *t* | *P>\|t\|* | *[0.025* | *0.975]* | *No. Observations* |
| --- | --- | --- | --- | --- | --- | --- | --- |
| Intercept | 0.4724 | 0.06 | 7.919 | <.001 | 0.344 | 0.601 | 15 |
| Occipital (Var) | -0.0093 | 0.06 | -0.156 | 0.879 | -0.138 | 0.12 | 15 |

**Table S11.**
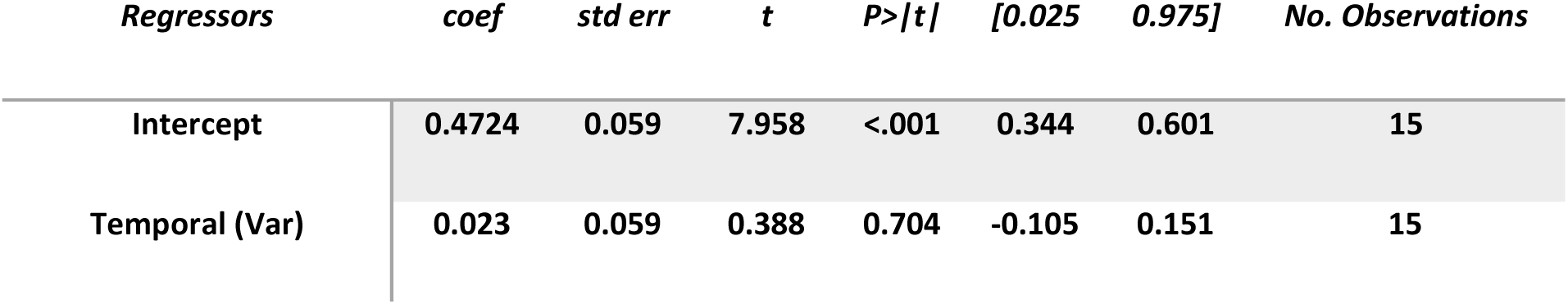
Details of statistical results in the relationship between beta of P(correct) and EEG beta of variance (Temporal).

**Table S12.** Details of statistical results in the relationship between beta of Fisher Information and EEG beta of variance (Centro-Parietal).

| *Regressors* | *coef* | *std err* | *t* | *P>\|t\|* | *[0.025* | *0.975]* | *No. Observations* |
| --- | --- | --- | --- | --- | --- | --- | --- |
| Intercept | 0.2209 | 0.062 | 3.578 | 0.003 | 0.088 | 0.354 | 15 |
| Centro-Parietal (Var) | 0.061 | 0.062 | 0.988 | 0.341 | -0.072 | 0.194 | 15 |

**Table S13.** Details of statistical results in the relationship between beta of Fisher Information and EEG beta of variance (Lateral-Parietal).

| *Regressors* | *coef* | *std err* | *t* | *P>\|t\|* | *[0.025* | *0.975]* | *No. Observations* |
| --- | --- | --- | --- | --- | --- | --- | --- |
| Intercept | 0.2209 | 0.064 | 3.462 | 0.004 | 0.083 | 0.359 | 15 |
| Lateral-Parietal (Var) | -0.0182 | 0.064 | -0.285 | 0.78 | -0.156 | 0.12 | 15 |

**Table S14.** Details of statistical results in the relationship between beta of Fisher Information and EEG beta of variance (Occipital).

| *Regressors* | *coef* | *std err* | *t* | *P>\|t\|* | *[0.025* | *0.975]* | *No. Observations* |
| --- | --- | --- | --- | --- | --- | --- | --- |
| Intercept | 0.2209 | 0.063 | 3.517 | 0.004 | 0.085 | 0.357 | 15 |
| Occipital (Var) | 0.0446 | 0.063 | 0.71 | 0.49 | -0.091 | 0.18 | 15 |

**Table S15.** Details of statistical results in the relationship between beta of Fisher Information and EEG beta of variance (Temporal).

| *Regressors* | *coef* | *std err* | *t* | *P>\|t\|* | *[0.025* | *0.975]* | *No. Observations* |
| --- | --- | --- | --- | --- | --- | --- | --- |
| Intercept | 0.2209 | 0.064 | 3.458 | 0.004 | 0.083 | 0.359 | 15 |
| Temporal (Var) | -0.0145 | 0.064 | -0.227 | 0.824 | -0.153 | 0.123 | 15 |

### Relation between AUC score and EEG beta of variance

We analyzed how the EEG coefficients of standard deviation and variance of stimulus from different regions impacted the AUC score by applying the same regression model for model’s parameters (Eq. S3).

**Table S16.** Details of statistical results in the relationship between AUC and EEG beta of variance and standard deviation.

| *Regressors* | *coef* | *std err* | *t* | *P>\|t\|* | *[0.025* | *0.975]* | No.<br>Observations |
| --- | --- | --- | --- | --- | --- | --- | --- |
| ***Intercept*** | **0.6353** | **0.01** | **64.498** | **<0.001** | **0.608** | **0.663** | **15** |
| ***Frontal (Var)*** | **0.2443** | **0.47** | **0.519** | **0.631** | **-1.062** | **1.55** | **15** |
| ***Lateral-Parietal (Var)*** | **0.7148** | **0.196** | **3.642** | **0.022** | **0.17** | **1.26** | **15** |
| ***Centro-Parietal (Var)*** | **1.3116** | **0.32** | **4.103** | **0.015** | **0.424** | **2.199** | **15** |
| ***Occipital (Var)*** | **-1.4448** | **0.417** | **-3.463** | **0.026** | **-2.603** | **-0.286** | **15** |
| ***Temporal (Var)*** | **-2.0135** | **0.531** | **-3.789** | **0.019** | **-3.489** | **-0.538** | **15** |
| ***Frontal (StdDev)*** | **0.0682** | **0.475** | **0.143** | **0.893** | **-1.251** | **1.387** | **15** |
| ***Lateral-Parietal (StdDev)*** | **0.741** | **0.209** | **3.545** | **0.024** | **0.161** | **1.321** | **15** |
| ***Centro-Parietal (StdDev)*** | **1.2081** | **0.289** | **4.177** | **0.014** | **0.405** | **2.011** | **15** |
| ***Occipital (StdDev)*** | **-1.2787** | **0.383** | **-3.34** | **0.029** | **-2.342** | **-0.216** | **15** |
| ***Temporal (StdDev)*** | **-1.978** | **0.527** | **-3.754** | **0.02** | **-3.441** | **-0.515** | **15** |

**Figure S12.**
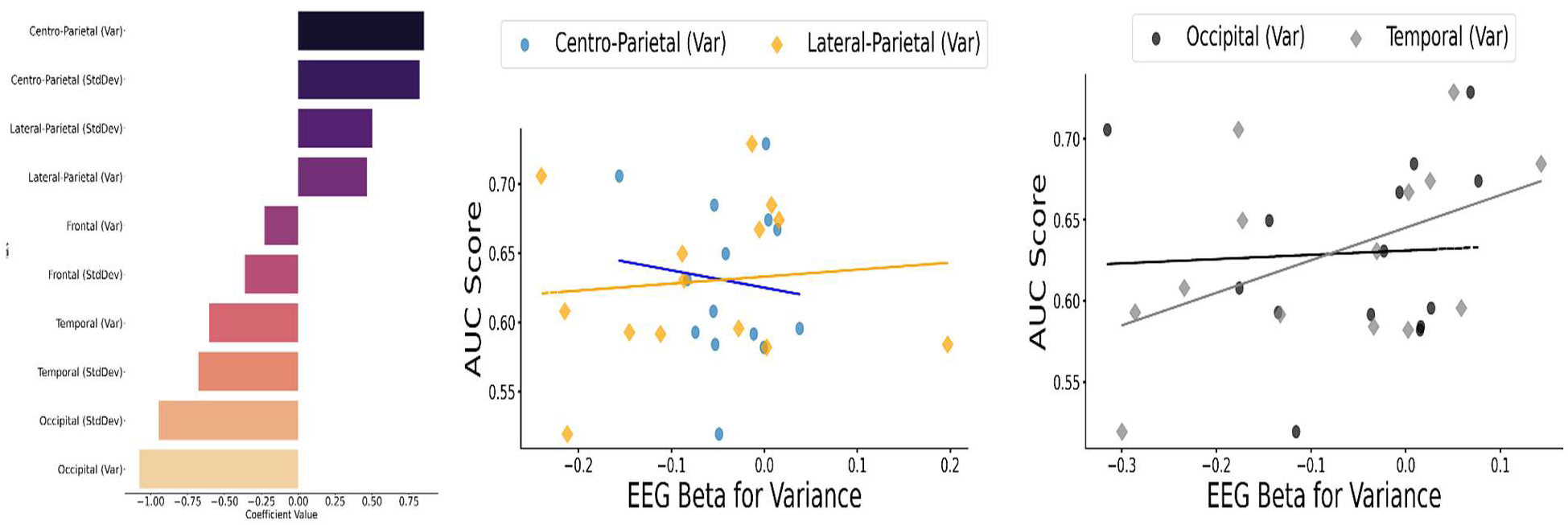
Exploring Neural Correlates of Variance and Model Parameters as Predictors of Metacognitive Sensitivity.

